# *ORM*-mediated regulation of sphingolipid biosynthesis is essential for nodule formation in *Aeschynomene evenia*

**DOI:** 10.1101/2023.09.12.557380

**Authors:** Nico Nouwen, Marjorie Pervent, Franck El M’Chirgui, Frédérique Tellier, Maëlle Rios, Natasha Horta Araújo, Christophe Klopp, Frédéric Gressent, Jean-François Arrighi

**Affiliations:** IRD, Plant Health Institute of Montpellier (PHIM), UMR IRD/SupAgro/INRAE/UM/CIRAD, TA-A82/J - Campus de Baillarguet, 34398 Montpellier, France.; INRAE, Plant Health Institute of Montpellier (PHIM), UMR IRD/SupAgro/INRAE/UM/CIRAD, TA-A82/J - Campus de Baillarguet, 34398 Montpellier, France.; Université Paris-Saclay, INRAE, AgroParisTech, Institut Jean-Pierre Bourgin (IJPB), 78000, Versailles, France.; Plateforme bioinformatique Genotoul, BioinfoMics, UR875 Biométrie et Intelligence Artificielle, INRAE, Castanet-Tolosan, France.

## Abstract

Legumes are able to establish symbiotic interactions with nitrogen-fixing rhizobia that are accomodated in root-derived organs, the nodules. Rhizobia recognition triggers a plant symbiotic signalling pathway activating two coordinated processes: infection and nodule organogenesis. How these are orchestrated in legumes species utilizing intercellular infection and lateral root-base nodulation remain elusive. Here, we show that *Aeschynomene evenia* OROSOMUCOID PROTEIN 1 (AeORM1), a key regulator of sphingolipid biosynthesis, is required for nodule formation in this legume species. Using *A. evenia orm1* mutants, we demonstrate that alterations in AeORM1 function result in numerous early aborted nodules, exhibiting defense-like reactions, and shortened lateral roots. Consistantly, *AeORM1* was expressed during lateral root initiation and elongation, including at lateral root bases where nodule primordia form in the presence of symbiotic bradyrhizobia. Sphingolipidomics revealed that mutations in *AeORM1* leaded to sphingolipid overaccumulation in roots, in particular the very-long-chain-fatty-acid (VLCFA)-containing ceramids relative to the wild-type plants. Taken together, our findings reveal that AeORM1-regulated sphingolipid homeostasis is essential for rhizobial infection and nodule organogenesis, and indicate a shared requirement for nodule formation and lateral root development in *A. evenia*.

## Introduction

Legumes have the ability to establish a nitrogen-fixing symbiosis with bacteria collectively named rhizobia. They form specific root organs, the nodules, where rhizobia are housed to convert atmospheric dinitrogen into nitrogen organic compounds in exchange for supply of carbon sources. In most rhizobium-legume interactions, nodulation occurs in a susceptible root zone with developing root hairs. Compatible rhizobia colonize the root hair surface and induce their curling to entrap a microbial colony in an infection chamber from which an infection thread develops. This tubular structure guides rhizobia to the nodule primordium that is distantly formed in the root cortex and where they are released. Bacterial accommodation is accompanied with their differentiation into bacteroids that are dedicated to nitrogen fixation (Roy et al., 2020).

The molecular basis of the rhizobial symbiosis has been well studied in two temperate model legumes, *Medicago truncatula* and *Lotus japonicus*, in which many nodulation genes have been identified. This symbiosis hinges on the recognition of rhizobial Nod factors by specific plasma membrane-localized LysM-RLK receptors. In turn, this recognition triggers a Nod signaling pathway leading to the activation of a network of nuclear transcription factors that induce the expression of symbiotic genes orchestrating the infection process and nodule organogenesis. Among them, NIN was shown to play an important role in the transition of the nodule into a nitrogen-fixing state (Feng et al., 2021; Roy et al., 2020). These gene discoveries provided valuable information on the molecular mechanisms involved in rhizobial symbiosis, among which: i) part of the signaling and infection genes are also involved in the more ancient endomycorrhizal symbiosis (Gobbato et al., 2015), ii) the infection thread polarized growth involves a multiprotein infectosome complex linked to vesicle trafficking and the cytoskeletton (Lace et al., 2023; Liu et al., 2019), iii) several plant genes are required to prevent defense reactions in nodules (Berrabah et al., 2018) and iv) the developmental program of nodules overlpas with lateral roots (Schiessl et al., 2019; Soyano et al., 2019).

Although the above described symbiotic mechanisms are likely representative of what occurs in many legume species, the legume family is huge and diverse (∼20,000 species), and significant variations on the theme of nodulation has been described. As example, in *Aeschynomene* species, root nodules form exclusively at a lateral root base (LRB) where a rosette of axillary root hairs is present. Bradyrhizobia enter the root at the base of these axillary root hairs and subsequently progress intercellularly in the root cortex. Eventually, the bacteria are endocyted by a few cortical cells that start to divide repeatedly to give rise to the nodule (Bonaldi et al., 2011). LRB nodulation is also found in other legume species like peanut - *Arachis hypogaea* - and *Sesbania rostrata* (Sharma et al., 2020; Capoen et al., 2010), and in 25% of the legume genera rhizobial infection occurs via intercellular penetration with again peanut as most known example (Quilbé et al., 2022a). An even more rare variation is found in approximately 20 *Aeschynomene* species that interact symbiotically with photosynthetic *Bradyrhizobium* strains. In these species, symbiosis occurs without Nod factor recognition (Chaintreuil et al., 2018; Giraud et al., 2007). In that case, LRB nodulation-associated axillary root hairs and intercellular infection show specific adaptations that likely facilitate the so-called Nod-independent symbiosis (Bonaldi et al., 2011).

The recent release of the *Aeschynomene evenia* genome and a collection of EMS-induced nodulation mutants has positioned it as a valuable model plant to study these nodulation features (Quilbé et al., 2021, 2022a, 2022b). Such analysis is predicted to enable the identification of new symbiotic mechanisms and to complement the information on nodulation as obtained with the historical model legumes. Initial genetic studies using *A. evenia* mutants demonstrated the conservation of several genes of the symbiotic signaling pathway, *AePollux*, *AeCCaMK*, *AeCyclops*, *AeNSP2* and *AeNIN* but not of those coding for the upstream Nod factor receptors. The discovery of *AeCRK*, encoding a receptor-like kinase required for symbiosis, also presented an important avenue to further investigate how the Nod-independent symbiosis is activated (Quilbé et al., 2021). Further analyses revealed that these genes intervened during different steps of intercellular infection, and that AeNSP2 has an additional function in controlling axillary root hair formation at lateral root bases, which constitute bradyrhizobia-colonized infection sites in *A. evenia* (Quilbé et al., 2022b).

Thus, although progress has been made in the identification of genes required for the activation of Nod-independent symbiotic and intercellular infection in *A. evenia*, our mechanistic understanding of intercellular infection and LRB nodulation remains in its infancy. In this study, we reported the mutant-based identification and functional characterization of the *AeORM1* gene predicted to encode an orosomucoid (ORM) protein, whose Arabidopsis orthologs are negative regulators of sphingolipid synthesis. In accordance with a gene expression during nodule and lateral root formation, alteration of ORM function resulted in shorter lateral roots and, in the presence of *Bradyrhizobium*, in early nodule abortion accompanied with defense-like responses. We also demonstrated that ORM mutations leaded to significant modifications in sphingolipid composition in roots. These findings strongly suggest that ORM regulation of sphingolipid homeostasis plays a key role during nodule formation and further links nodule development to lateral root development in *A. evenia*.

## Results

### *A. evenia ORM1* gene is required for rhizobial symbiosis

To uncover the molecular mechanisms underpinning the original nodulation properties found in *A. evenia*, we recently screened an ethyl-methane-sulfonate (EMS) mutagenized population for defects in nodule formation with the photosynthetic *Bradyrhizobium* strain ORS278 (Quilbé et al., 2021). Three nodulation mutants named P35, Q33 and AG2 were isolated based on nitrogen starvation symptoms (i.e. under-developed plants with yellowing leaves), that were simiar to those observed for Nod^-^ (no nodule) mutants such as *ccamk-3* (Figure 1A). In contrast to the WT line that produced pink-colored nodules and the completely noduleless *ccamk-3* mutant, the roots of P35, Q33 and AG2 mutant plants contained numerous brown spots at the base of lateral roots (Figure 1B). While the base of lateral roots of the *ccamk-3* mutant exhibited only the typical orange crowns of axillary root hairs, in the three newly analyzed mutants, the brown spots were round-shaped bumps, suggesting early abortion of nodule formation associated with defense-like pigment accumulation (Figure 1B – zooms).

**Figure 1.**
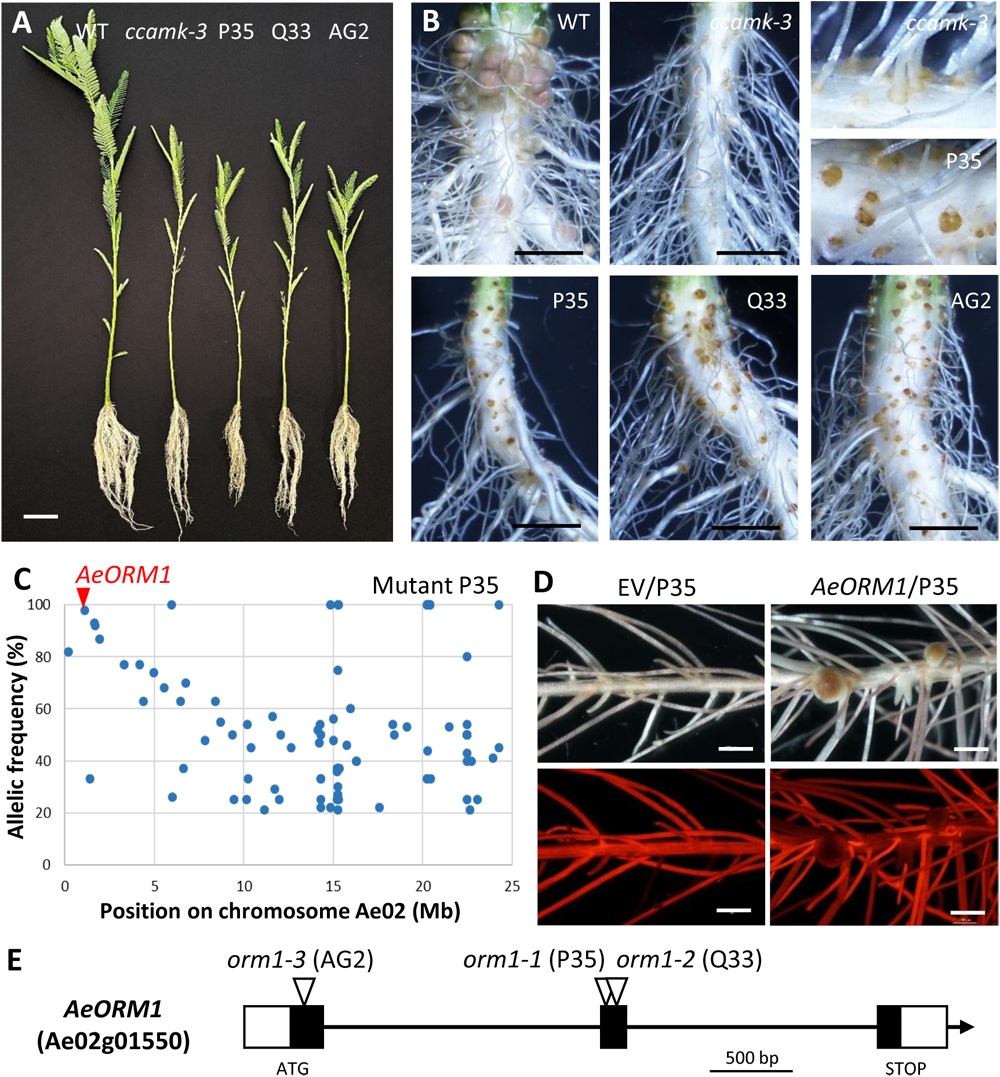
Mutations in *AeORM1* disrupt nodule formation and cause defense-like reactions. **(A)** Growth phenotype of WT, *ccamk-3*, and P35, Q33 and AG2 mutant plants grown inoculated with *Bradyrhizobium* strain ORS278 and grown for 28 days in greenhouse conditions. Bar = 5 cm. (**B**) Symbiotic phenotype of roots from plants shown in panel (A). Root of the WT carries pink nodules, while *ccamk-3* root is completely noduleless and the P35, Q33 and AG2 mutat roots display numerous brown spots at nodulation sites. Upper right panel show a zoom of nodulation sites on *ccamk-3* and the P35 mutant roots. Bars = 5 mm. **(B)** Frequency of the EMS-induced mutant alleles in bulks of backcrossed F_2_ mutant plants of the P35 mutant as obtained by the Mapping-by-Sequencing approach. The SNP representing the putative causal mutation is indicated by the red arrow head (AF = 98%). (**D**) Functional complementation assays. Transgenic hairy roots of *orm1-1* mutant plant containing the empty vector (EV) remain noduleless, whereas hairy roots containing the ProAeORM1-AeORM1 construct contain nodules at 28 dpi. Upper panels, bright field images of hairy roots and the lower panels, epifluorescent microscopic images showing DsRed expression in the same transgenic roots. Bars = 2 mm. (**E**) Structure of the *AeORM1* gene and positions of the EMS mutations identified in the P35 (*orm1-1*), Q33 (*orm1-2*) and AG2 (*orm1-3*) mutants. White boxes correspond to UTR regions, black boxes to exons and arrow heads indicate locations of the mutations.

F_2_ progenies generated from crosses with the WT line segregated in a 3 :1 ratio of plants with pink nodules to plants with only brown spots, suggesting that each mutation is monogenic and recessive (Supplemental Table 1). To identify the mutations causing this nodulation phenotype, we conducted a Mapping-by-Sequencing approach by sequencing pooled DNAs from mutant plants within the segregating F_2_ populations. For the three mutants, a genetic linkage was identified at the same location near the top of the Ae02 chromosome (Figure 1C, Supplemental Figure 1). Closer inspection of this region revealed the presence of distinct mutations in the Ae02g01550 gene with a mutant allelic frequency of 98, 98 and 82% for the P35, Q33 and AG2 mutants, respectively (Supplemental Table 1). Functional annotation of the Ae02g01550 gene indicated that it is predicted to encode an orosomucoid (ORM) protein. ORM proteins are known to be localized in the endoplasmic reticulum (ER) where they act as major negative regulators of sphingolipid biosynthesis in plants and other eukaryotes (Kimberlin et al., 2016 ; Li et al., 2016). Therefore, we named this candidate gene *AeORM1*. To validate the *AeORM1* gene, allelism tests were performed by crossing the three mutants between them (Supplemental Table 2). All the F_1_ plants produced brown spots after inoculation with *Bradyrhizobium* ORS278 strain. In addition, the full-length WT CDS was cloned with its native promoter region (i.e. ∼ 2.1 kb upstream of the predicted start codon) and expressed using the hairy root transformation protocol in roots of P35 plants. In contrast to the P35 plants transformed with the empty vector that were completely noduleless, nodules readily developed on the complemented root systems after inoculation with *Bradyrhizobium* ORS278 (Figure 1D, Supplemental Table 3). Taken together, these results unambiguously indicate that the mutations in *AeORM1* are responsible for the nodulation phenotype. Accordingly, we designated the alleles in the P35, Q33 and AG2 mutants *orm1-1*, *orm1-2* and *orm1-3*, respectively. In the three exon-containing *AeORM1* gene, *orm1-3* mutation falls in exon 1 while the *orm1-1* and *orm1-2* mutations are located in exon 2 (Figure 1E, Supplemental Table 1).

### *AeORM1* belongs to a small family of highly conserved genes

To our knowledge, the involvement of *ORM* genes in rhizobial symbiosis has not yet been reported. This prompted us to characterize the *ORM* gene family in legumes by searching for *AeORM1* homologs in 11 Papilionoideae and 2 Caesalpionoideae species for which the genome has been completely sequenced. Arabidopsis and rice *ORM* genes were included for comparison. One to four *ORM* homolog genes were retrieved for each considered species. Phylogenetic analysis showed that Arabidopsis and rice *ORM* genes form two independent clades while legume *ORM* genes are organized in three clusters (Figure 2A). One contained Caesalpinoideae genes and the two others had Papilionoideae genes. Interestingly, *AeORM1* clusters with *ORM1* representatives from all other Papilionoideae, forming the ORM1 clade.

**Figure 2.**
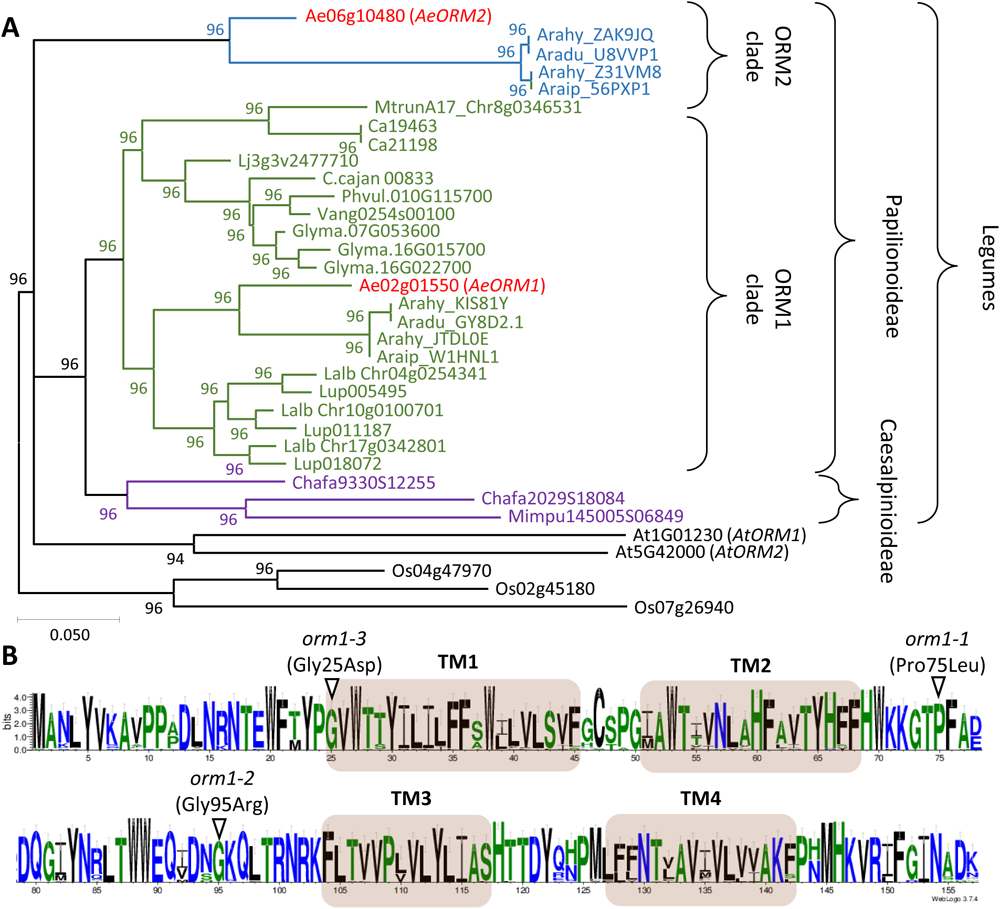
*AeORM1* is part of a highly conserved gene family. (**A**) Maximum likelihood tree of plant ORM genes based on MUSCLE nucleotidic alignment with 1000 bootstrap values. Selected species are: *Aeschynomene evenia* (Ae), *Arabidopsis thaliana* (AT), *Arachis hypogaea* (Arahy), *Arachis duranensis* (Aradu), *Arachis ipaiensis* (Araip), *Cajanus cajan* (C. cajan), *Chamaecrista fasciculata* (Chafa*), Cicer arietum* (Ca), *Glycine max* (Glyma), *Lupinus albus* (Lalb), *Lupinus angustifolius* (Lup), *Lotus japonicus* (Lj), *Medicago truncatula* (Mtrun), *Mimosa pudica* (Mimpu), *Oryza sativa* (Os), *Phaseolus vulgaris* (Phvul), *Vigna angularis* (Vang). Within the legume family, Caesalpinioideae genes are in purple and Papilionoideae genes are in green and blue to hightlight the presence of the ORM1 and ORM2 clades, respectively. (**B**) Sequence logo for plant ORM proteins generated after a MUSCLE alignment of the predicted protein sequences of genes used in (A) using the Weblogo software and showing the conserved amino acid (AA) residues. Blue: hydrophile AAs, green: Neutral AAs, black: hybrophobic AAs. Predicted transmembranes (TM) are indicated by grey boxes and AA changes resulting from the EMS mutations in *AeORM1* are indicated by arrow heads.

*A. evenia* contains a second gene copy of *ORM* gene that we named *AeORM2*. This copy has only counterparts in *Arachis* spp. with whom it forms a seperate cluster which we named the ORM2 clade. Synteny analysis confirmed orthologous and paralogous relationships of *ORM* genes among Papilionoideae species, reinforcing the idea that the ORM1 and ORM2 clades result from the polyploidy event at the base of Papilionoideae (Supplemental Figure 2) (Li et al., 2013). So, while only Dalbergioid legumes seem to have retained the duplicated ORM copies (such as *A. evenia*), others have only preserved the ORM1 copy (such as *M. truncatula* and *Lotus japonicus*). More recent gene or whole genome duplications are likely responsible for variations in gene copy numbers as found in different species (e. g. *Glycine max* and *Arachis hypogaea*).

Sequence alignment of the predicted ORM proteins revealed that they are almost all 157 amino acid (AA)-long (Supplemental Figure 3). Another striking feature was that these ORM proteins are highly conserved in their sequence throughout the plant kingdom. Indeed, AeORM1 and AeORM2 share 89.2% identity while AeORM1 shares 93% identity with the *M. truncatula* ORM protein and still an 84.1% identity with the more distantly related rice ORM proteins. Similar to Arabidopsis ORM proteins, legume ORM proteins are predicted to have four transmembrane domains, typically containing hydrophic and neutral AAs while remaining sequences are enriched in hydrophile AAs (Figure 2B). Interestingly, the mutations identified in the three *A. evenia orm1* mutants lead to AA substitutions in conserved residues of the first transmembrane domain (*orm1-3*: Gly25Asp) and the central loop (*orm1-1*: Pro75Leu and *orm1-2*: Gly95Arg) (Figure 2B). The drastically altered nodulation phenotype of the *orm1* mutants strongly indicate that these AA substitutions are detrimental for ORM activity.

### *AeORM1* expression associates with lateral root and nodule development

To get information on the potential involvements of the two *ORM* genes in *A. evenia*, their transcript levels were analyzed in the different plant organs and during symbiotic interactions. Both genes were found to be expressed ubiquitously in stems, leaves, flowers, pods, roots and during nodulation, based on the *A. evenia* Gene Atlas (Supplemental Figure 4A). *AeORM1* expression levels appeared to be ∼ 30% higher than those of *AeORM2* in all conditions, except in flowers. To confirm and extend the *A. evenia* Gene Atlas data, RT-qPCR analysis was conducted on RNA isolated from WT roots inoculated or not with *Bradyrhizobium* strain ORS278 (for nodulation) and *Rhizophagus irregularis* (for mycorrhization). Similar expression patterns were observed for both genes during the two symbioses (Supplemental Figure 4B). These data suggested that *AeORM1* and *AeORM2* might act in concert and that *AeORM1*, in addition to nodulation, may have other roles in the development of *A. evenia* plants.

To analyze the spatial expression of the *AeORM1* gene, we fused the ∼ 2.1 kb promoter region to the GUS reporter gene. This p*AeORM1*-*GUS* construct was used to transform WT *A. evenia* roots with *A. rhizogenes* and GUS activity was monitored in non-and inoculated roots. In non-inoculated roots, GUS staining was observed in lateral root primordia as found, in particular, in the younger part of the primary root (Figure 3A to C). Once emerged from the primary root, GUS staining was observed both at the base and tip of lateral roots (Figure 3D to F). After inoculation with *Bradyrhizobium* ORS278, enhanced GUS staining was observed at nodule primordium initiation sites located at lateral root bases and where both rhizobial infection and cell divisions occur (Figure 3G to K). When young nodules became apparent (i.e., at 4 dpi), GUS staining was observed in the region that surrounded the central infected tissue (Figure 3L). GUS staining persisted in mature nodules (i.e., at 11 dpi). However, sectioning and examination of these nodules by light microscopy revealed specific expression of p*AeORM1*-*GUS* at the nodule base and in a few cell layers encircling the central nitrogen-fixation zone (Figure 3M to O). Prolonged GUS staining of sectioned nodules never led to blue coloration of the central infected tissue.

**Figure 3.**
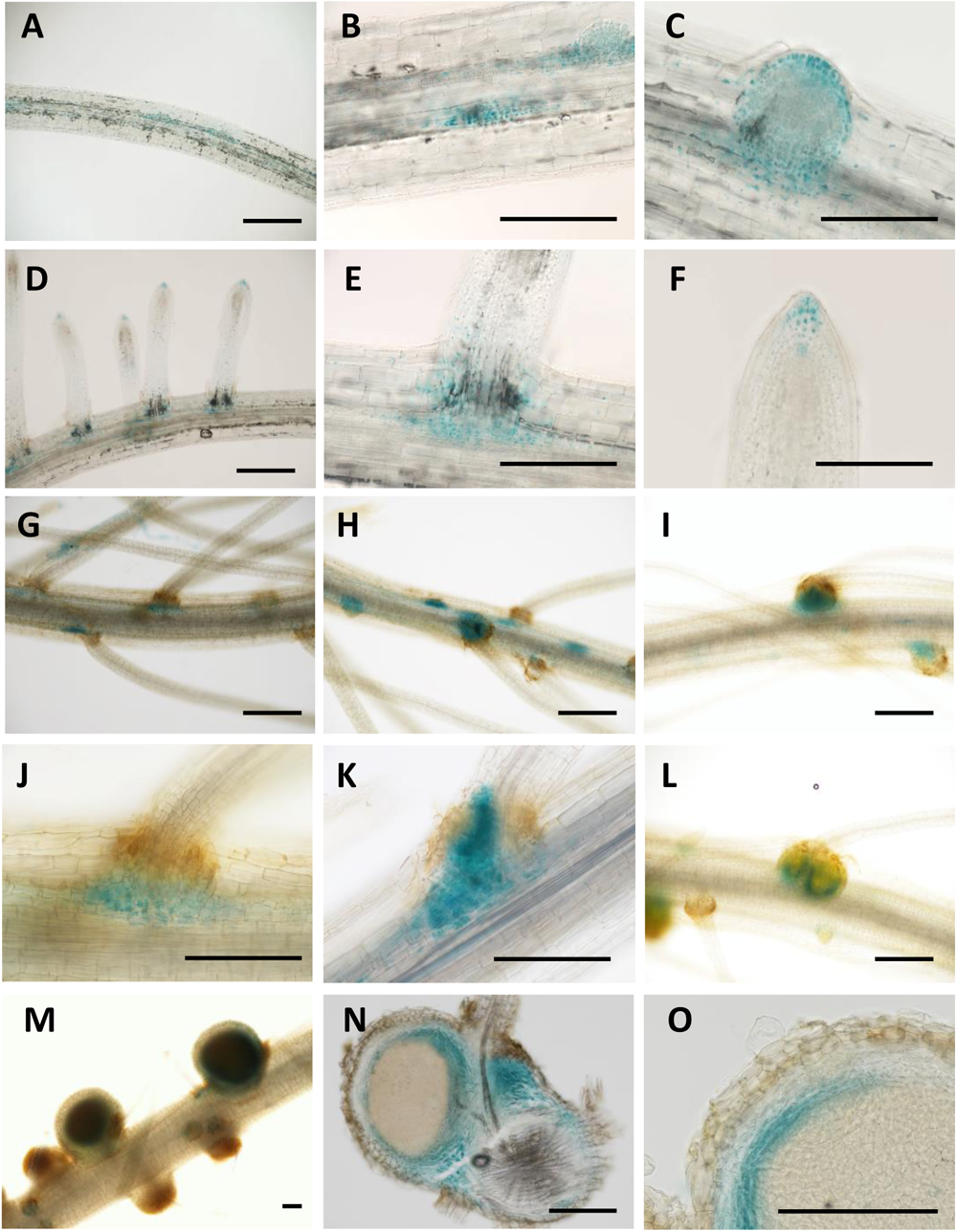
*AeORM1* is expressed during root and nodule initiation and development. Histochemical localization of GUS activity in *A. rhizogenes* roots of WT *A. evenia* transformed with proAeORM1-GUS. (**A**) to (**F**) Non-inoculated roots showing glucoronidase activity during lateral root initiation and development. (**A**) whole primary root, (**B**) and (**C**) zoom on inner and emerging root primordia, (**D**) whole primary root with elongating lateral roots, (**E**) and (**D**) base and apex of an elongating lateral root, respectively. Bars = 500 µm. (**G**) to (**L**) Glucoronidase activity in roots 4 days after inoculation with B*radyrhizobium* ORS278. The sequential enlargement of the zone containing glucoronidase activity is shown in (**G**), (**H**) and (**I**) respectively, and zooms showing (**J**) basal activity, (**K**) nodule primordium-associated activity and (**L**) activity relocalization in young nodules. Bars = 500 µm. (**M**) to (**O**) Glucoronidase activity in nodules at 11dpi. (**M**) Whole nodules showing intense blue coloration after staining roots with X-gluc. (**N**) Glucoronidase activity in a nodule section. (**O**) Zoom in a nodule section showing glucoronidase activity in only plant cells at the periphery of the central infected tissue. Bars = 200 µm.

### Nodule formation is compromised in *orm1* mutants

To associate the *AeORM1* expression pattern during nodulation with the observed mutant phenotypes, we studied the nodulation kinetics of the three *orm1* mutants after inoculation with *Bradyrhizobium* strain ORS278 and analyzed their symbiotic alterations by light microscopy at different time points (7, 10, 14 and 21 dpi). In WT plants, mature nodules became slightly pink at 7 dpi and this pink coloration intensified at later stages, an indication of leghemoglogin accumulation (Supplemental Figure 5). In contrast, no nodules developed in the three *orm1* mutants. However macroscopic examination of root sections evidenced limited cell divisions at the base of some lateral roots, indicative of nodule initiation, and brown spots, likely a result of accumulation of defense-like pigments. These features were discrete and visible as soon as 7 dpi but the presence of both bumps and brown spots was more pronounced at later time points (Supplemental Figure 5). A more detailed analysis was performed using a GUS-tagged version of strain ORS78. At 10 dpi, examination of whole and sectioned roots showed that the brown spot-containing bumps were lobe-shaped like during nodule formation in WT *A. evenia* (Arrighi et al., 2012), but a gradation in the development was observed in the different *orm1* mutant plants: with the *orm1-1* plants only limited cell divisions where observed whereas *orm1-3* plants contained more round-shaped bumps (Figure 4A). X-Gluc staining revealed that all three *orm1* mutants were colonized by bradyrhizobia. In this aspect, the *orm1-1* mutant was again the most severely affected with only a few inner infection pockets while in *orm1-2* and *orm1-3* mutant plants bumps with a central infected tissue were visible (Figure 4A). Dark brown spots were located in the vicinity of infection pockets in the *orm1-1* mutant plants and they were present in the form of more extended brownish areas in the outer and/or inner nodule tissues of both other *orm1* mutants (Figure 4A).

**Figure 4.**
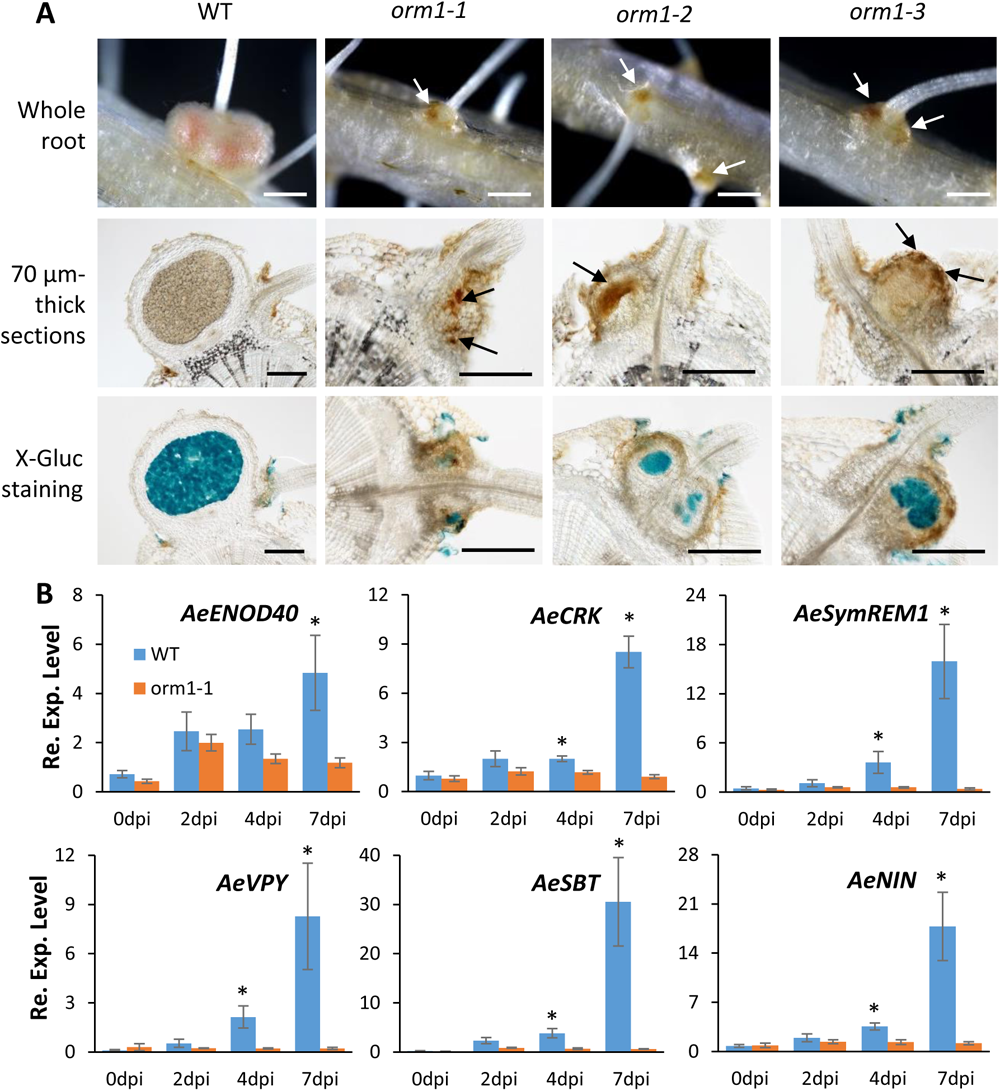
Mutations in *AeORM1* impair nodule development and alter symbiotic signaling. **(A)** Nodule development in WT and *orm1* mutant plants at 10 dpi with *Bradyrhizobium* ORS278-GUS. Whole roots (upper panels), 70-µm-thick root sections before and after X-Gluc staining (middle and lower panels, respectively). Arrows point to the brown pigments visible in sections of *orm1* mutant roots. Bars = 250 µm. (**B**) Expression of early nodulation genes in WT and *orm1-1* mutant plants. Expression of *AeENOD40*, *AeCRK*, *AeSymREM1*, *AeVPY*, *AeSBT*, and *AeNIN* in plant roots, was determined during nodulation kinetics with Bradyrhizobium ORS278 at 0, 2, 4 and 7-dpi by RT-qPCR analysis. Expression values were normalized using *AeEF1a* and *Ubiquitin* expression levels as standard. Means and standard deviations (s.d.) were derived from four biological replicates and asterisks above the bars represent statistically significant differences (Mann-Whitney test ; *P < 0,05).

Next, we examined the effect of mutation in *AeORM1* on the expression of several genes that were previously shown to be induced during the initiation of rhizobial symbiosis in *A. evenia*: *AeENOD40*, *AeCRK*, *AeSymREM1*, *AeVPY*, *AeSBT* and *AeNIN* (Quilbé et al., 2022). RT-qPCR analysis was used to assay their induction by *Bradyrhizobium* ORS278 in the *orm1-1* mutant that shows the earliest block in nodule formation. The expression of all analyzed genes is induced as soon as 2dpi and show a continuous increase in expression up to 7 dpi in WT plants (Figure 4B). In contrast, in the *orm1-1* mutant, *AeENOD40* expression reached a maximum at 2dpi and then decreased in the following points (4-7 dpi). A low induction of expression in early time points (2-4 dpi) and premature decrease in later time points (4-7 dpi) was observed for the *AeCRK*, *AeSymREM1*, *AeSBT* and *AeNIN* genes (Figure 4B). These altered expression patterns are consistent with the phenotypic symbiotic responses observed in the *orm1-1* mutant (i.e., limited cell divisions and infection). In contrast to the other anlaysed genes, the induction of the *AeVPY* gene was completely impaired in the *orm1-1* mutant (Figure 4B).

### Defense-like reactions and altered rhizobial accomodation in *orm1* mutants

Since all three *orm1* mutants produce discrete bumps with brown pigmentation after infection with *Bradyrhizobium* ORS278, we wanted to characterize this pigmentation in more detail. For this, we made use of roots inoculated with *Bradyhrizobium* strain ORS278-GUS and that at 14 dpi were sectioned and stained with X-Gluc before microscopic observation. Brown pigmentation was visible within the bumps of all three *orm1* mutants and also dark brown inner spots were observed in the *orm1-1* mutant (Figure 5A). Fluorescent micropscopic analysis showed that the brown areas had a green fluorescence when using a green fluorescent protein (GFP) filter but not the dark brown inner spots present in the *orm1-1* mutant (Figure 5B). Red fluorescent areas were also observed when using the mCherry filter and athough they showed some overlap with the green fluorescent areas, the red fluorescent areas were in general more extended within the bumps, suggesting that the brown pigmentations may correspond to different phenolic compounds (Figure 5C). The presence of phenolic compounds in *orm1* bumps was confirmed using staining root sections with potassium permanganate/methylene blue. Precipitates of blue stain at sites of brown pigmentation were observed in all three *orm1* mutants and were completely absent in sections of the WT nodules (Figure 5D). Finally, 3,3′-Diaminobenzidine (DAB) staining revealed higher concentrations of H_2_O_2_ in regions where brown pigments accumulated in the *orm1* bumps, suggesting a concommitant alteration of redox status in these areas (Figure 5E).

**Figure 5.**
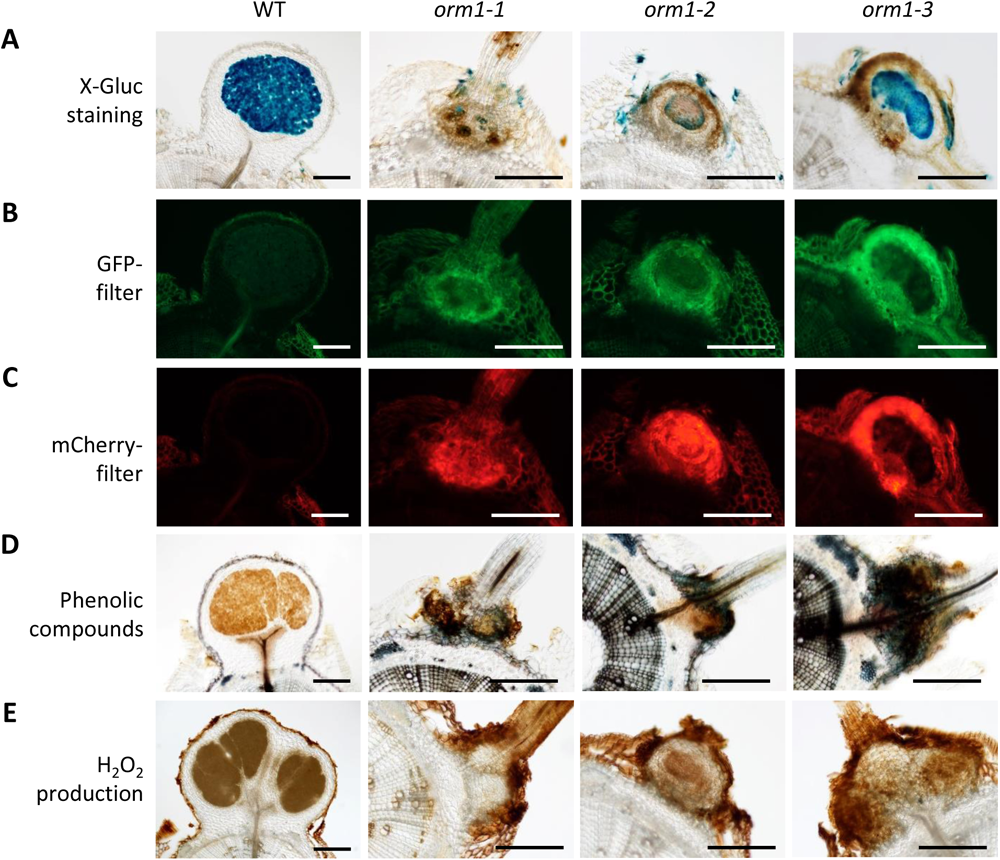
Mutations in *AeORM1* cause defense-like reactions. (**A**) to (**E**) 70 µm-thick root or nodule sections of WT and *orm1* mutant plants at 14 dpi with *Bradyrhizobium* ORS278 or ORS278-GUS strains. Bars = 250 µm. (**A**) Root sections of plant inoculated with the *Bradyrhizobium* ORS278-GUS strain after staining with X-gluc (the blue precipitate indicate the presence of bacteria). Note the presence of brown pigmentation in the root sections of the *orm1* mutant plants. (**B**) Fluorescent images of nodule sections shown in (**A**) using a GFP filter. In *orm1* mutants, green fluorescence co-localizes with the pale brown regions. Note : for all images, the same exposure time was used. (**C**) Fluorescent images of nodule sections shown in (**A**) using a mCherry filter. In *orm1* mutants, the red fluorescence co-localizes with both pale brown regions and intense dark brown spots. (**D**) Staining of nodule sections of plants inoculated with *Bradyrhizobium* ORS278 with potassium permanganate (KMnO_4_) and methylene blue. Blue staining indicates the presence of phenolic compounds in the plant tissue. (**E**) Staining of nodule sections of plants inoculated with *Bradyrhizobium* ORS278 with 3’,3’ diaminobenzidine (3’,3’ DAB) to detect hydrogen peroxyde (H_2_O_2_) production.

To determine to which extent the observed defense-like reactions are accompanied with defects in rhizobial colonization of symbiotic cells and rhizobial differentiation into bacteroids, *orm1* mutants were inoculated with a GFP-tagged version of the *Bradyrhizobium* strain ORS278 and at 14 dpi root sections were observed by confocal microscopy. Analysis of WT nodule sections revealed that the central tissue was largely occupied by infected cells packed with green fluorescent bacteria (Figure 6A). In contrast, *orm1* bumps were characterized by infected plant cells that were unevenly filled and that contained less bacteria than the WT. Strong autofluorescence was visible both within and outside of the central tissue (Figure 6A). Mature nodules of the WT plants contained typical spherical bacteroids while only elongated bacteria were observed in bumps of the three *orm1* mutants, suggesting an impaired differentiation process (Figure 6B). The block in bradyrhizobia differentiation correlated with the nitrogen-starvation symptoms of the plants, the early arrest in nodule development resulting in bumps that were hardly visible by the naked eye and the absence of nitrogenase enzyme activity in acetylene reduction assays (Figure 6C-E).

**Figure 6.**
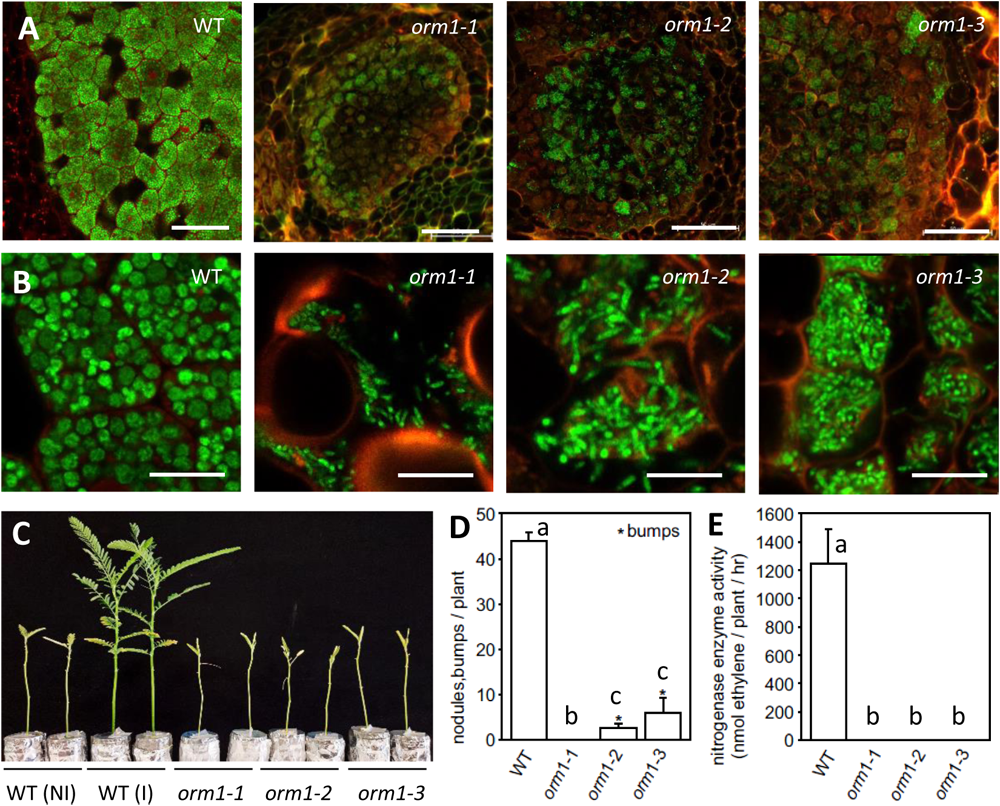
Mutations in *AeORM1* impair bacterial differentiation and nodule functionning. (**A**) Confocal microscopic analysis of longitudinal 70 µm-thick sections of nodules of plants inoculated with *Bradyrhizobium* ORS278-GFP at 14 dpi. Plant cell walls are visualised using their autofluorescent characteristics. Bars = 50 µM. (**B**) High magnification images of nodule sections showing the bacterial cell morphology in infected plant cells. At 14 dpi, bacteria are spherical in nodule cells of WT plants and elongated in infected cells of the *orm1* mutant plants. Bars = 10 µm. (**C**) Growth of non-inoculated (NI) and inoculated (I) plants (aerial part) after cultivation under hydroponic condition in a growth chamber. Images were taken 21 days after inoculation with *Bradyrhizobium* ORS278. (**D**,**E**) Number of bumps and nodules on plants (**D**) and nitrogenase enzyme activity measured by the acetylene reduction assay (**E**) of plants at 21 dpi. Error bars represent s.d. (n=6). Different letters represent significant differences determined using a Pairwise Wilcoxon test, P < 0.05.

#### *AeORM1* is not essential for mycorrhization

As many genes important for rhizobial symbiosis are also involved in mycorrhizal symbiosis, we investigated the role of *AeORM1* in the formation of arbuscular mycorrhiza (AM). Root phenotyping, 6 weeks after inoculation with spores of *Rhizophagus irregularis*, revealed that similar as to the WT plants, roots of the three *orm1* mutants contained fungal hyphae, arbuscules and vesicles being (Figure 7A). To refine the analysis, we focused on the the strongest allele mutant, *orm1-1*, and quantified the AM colonization. At 6 wpi, the mycorrhization frequency and intensity of the mutant were slightly higher compared to the WT, suggesting that *orm1* mutants are normally mycorrhized (Figure 7B and C). To further characterize AM symbiosis in *orm1* mutants, the expression of the marker genes *AeRAM1*, *AeVPY*, *AeSTR*, *AeSBTM1* were next investigated by RT-qPCR. These plant genes were previously shown to be strongly induced by AM infection in *A. evenia* (Quilbé et al., 2022). In the *orm1-1* mutant, the induction levels of all tested genes were similar to those in WT plants (Figure 7D). Quantification of the presence of the *RiLSU* and *RiGADPH* genes (markers of fungal biomass) in the root tissue, indicated that the abundancy of *R. irregularis* in *orm1-1* roots was equal to the WT roots (Figure 7D). All above results indicate that *AeORM1* is not essential for mycorrhization.

**Figure 7.**
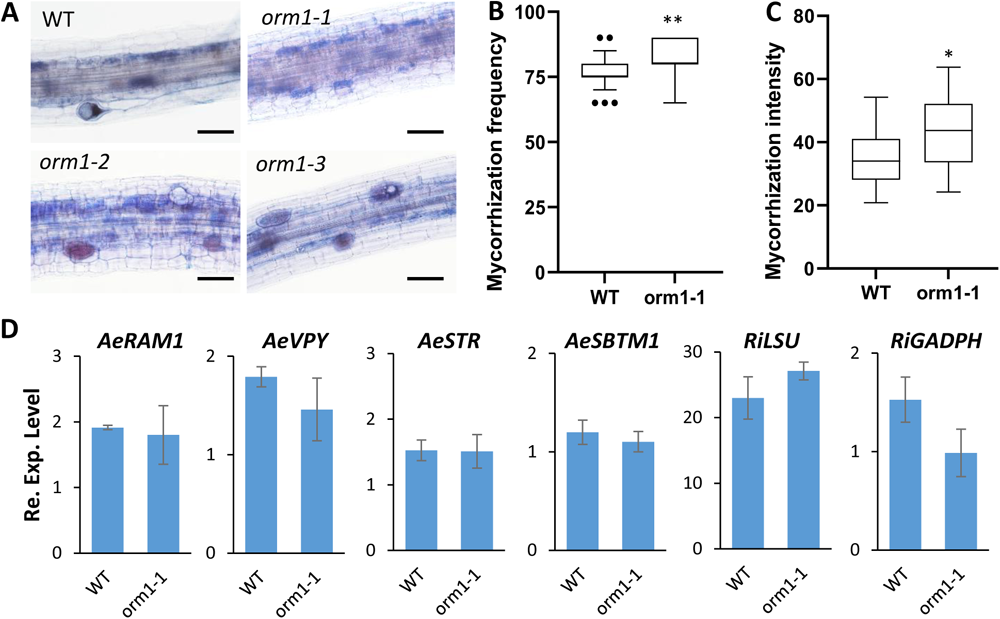
*AeORM1* does not play an essential role in mycorrhization. (**A**) Representative images of fungal colonization patterns observed in WT and *orm1* mutant plants. Bars = 50 µm. (**B**) Mycorrhization frequency and (**C**) intensity both expressed in % WT and *orm1-1* mutant plants. Box plots show median (central segment), second to third quartiles (box), minimum and maximum ranges (whiskers), and outliers (single points). Data are from four biological repeats, each with six plants/line. *P < 0.05, ** P < 0,001, significant differences between WT and *orm1-1* mutant plants using a Mann-Whitney test. (**D**) Expression of mycorrhization-induced genes in WT and *orm1-1* mutant plants. Expression of *AeRAM1*, *AeVPY*, *AeSTR*, *AeVPY*, *AeSBTM1*, and the two fungal genes *RiLSU* and *RiGADPH*, was determined by RT-qPCR using plant roots inoculated with *R. irregularis* and cultivated for 6 weeks. Expression values were normalized using *AeEF1a* and *Ubiquitin* expression levels as standard. Means and standard deviation (s.d.) were derived from three biological replicates. For each analysed gene, no statistically significant differences were observed between WT and *orm1-1* mutant plnats (Mann-Whitney test ; P < 0,05).

### *AeORM1* is important for normal lateral root development

*AeORM1* expression in non-symbiotic organs suggested that the role of this gene might not be restricted to rhizobial symbiosis. Although no detailed quantitative analysis was performed, in greenhouse conditions where plants are grown in compost complemented with organic fertilizers, all three allelic *orm1* mutants could produce seeds normally. This indicates that alterations in *AeORM1* have no major effect on the aerial plant development. In contrast, when analyzing the symbiotic phenotype for the three allelic *orm1* lines using hydroponic growth conditions in growth chamber, we noticed that young mutant plants have shorter lateral roots as compared to the WT plants. To avoid potential influences of age and nitrogen fixation on lateral root development, we assessed the root system architecture of the WT line and the three *orm1* mutants, cultured in liquid BNM medium containing 0.5 mM KNO_3_ for 10 days and without inoculation. Two out of the three *orm1* mutants had primary roots that were slightly shorter as compared to the WT plants while the lateral root number was found to be globally the same in all analyzed lines (Figure 8A and B). In contrast, significant variations in the lateral root length were found in all mutant lines relative to the WT, with the *orm1-1* mutant showing the most altered root phenotype (Figure 8C and D). Thus, *orm1* mutants have both a nodulation phenotype and an alteration in lateral root elongation. This suggests that *AeORM1* is involved in both lateral root and nodule development, in accordance with the observed expression pattern of this gene.

**Figure 8.**
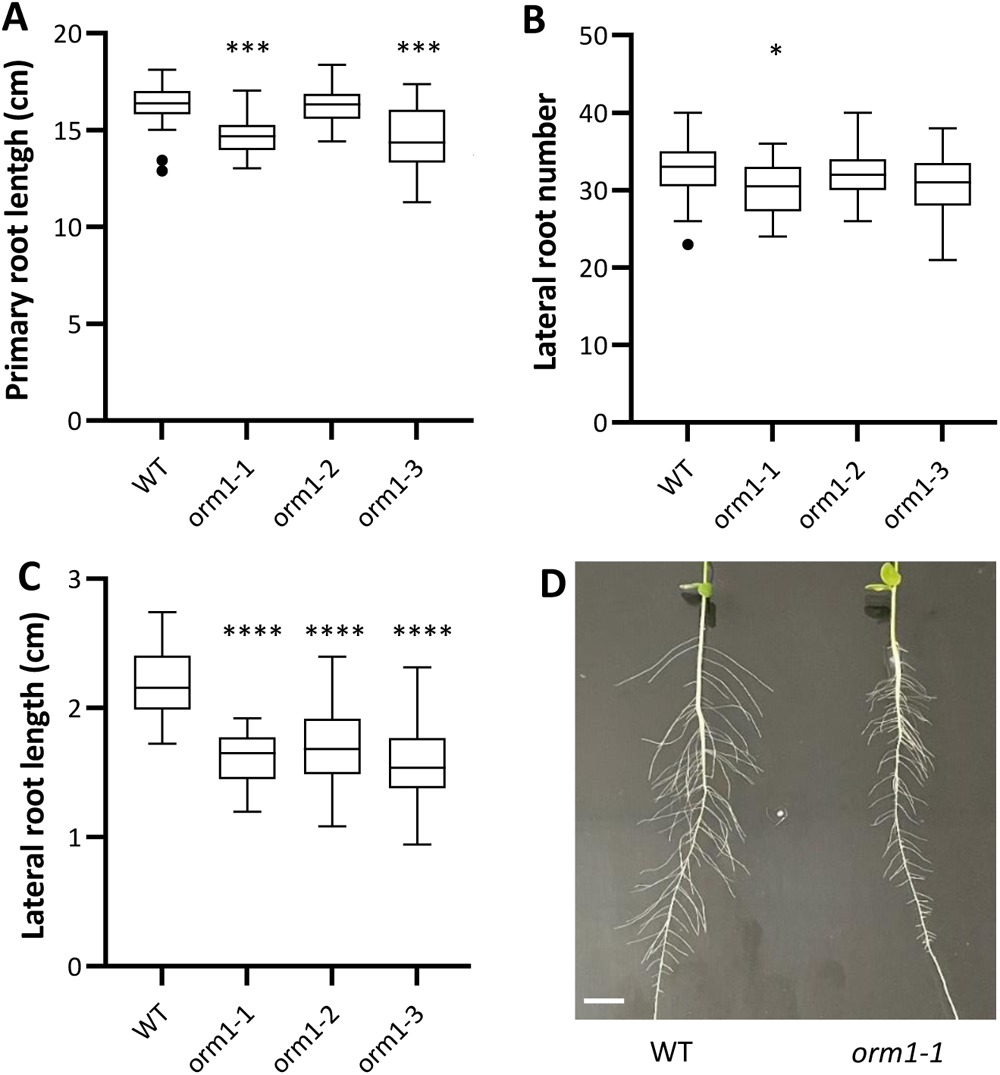
Lateral Root Development is Reduced in *orm1* Mutants. Analysis of the root system architecture of 10-day-old *A. evenia* wild-type (WT) and *orm1* mutant plants. (**A**) to (**C**) Box plots showing the median (central segment), second to third quartiles (box), minimum and maximum ranges (whiskers), and outliers (single points) of measurement of (**A**) primary root length; (**B**) lateral root number on a segment of 3 cm in the upper part of the primary root and (**C**) lateral root length on the same segement of 3 cm as in (**B**). *P < 0.05, ***P < 0.001 and ****P < 0.0001, significant differences between WT plants and each *orm1* mutant using a one-way Kruskal-Wallis test. n = 21 (WT), 24 (*orm1-1*), 24 (*orm1-2*) and 17 (*orm1-3*). (**D**) Images showing a primary root of a WT and *orm1-1* plant with emerging lateral roots. Bars = 1 cm.

### Mutations in *AeORM1* alter sphingolipid contents in roots

ORM proteins act as major negative regulators of the first step of the sphingolipid biosynthetic pathway where the condensation of Ser with palmitoyl-CoA give rise to long-chain bases (LCBs). These LCBs can be further modified and paired with structurally diverse fatty acids to produce ceramids (CER) and hydroxyceramides (hCER) that provide the backbone for more complex sphingolipids, including glucosylceramides (GlcCER) and different glycosylinositolphosphoceramides (GIPC) (Alsiyabi et al., 2021). To investigate the contribution of *AeORM1* in sphingolipid biosynthesis, we first analyzed the major classes of sphingolipids in the roots of WT plants and compared roots grown in BNM medium containing either 0.5 or 5 mM KNO_3_. Sphingolipid quantification on 4-week-old WT roots revealed little to no differences between the low and high nitrogen conditions (Figure 9A-E), indicating that the plant nitrogen status has no major impact on the root sphingolipid composition.

**Figure 9.**
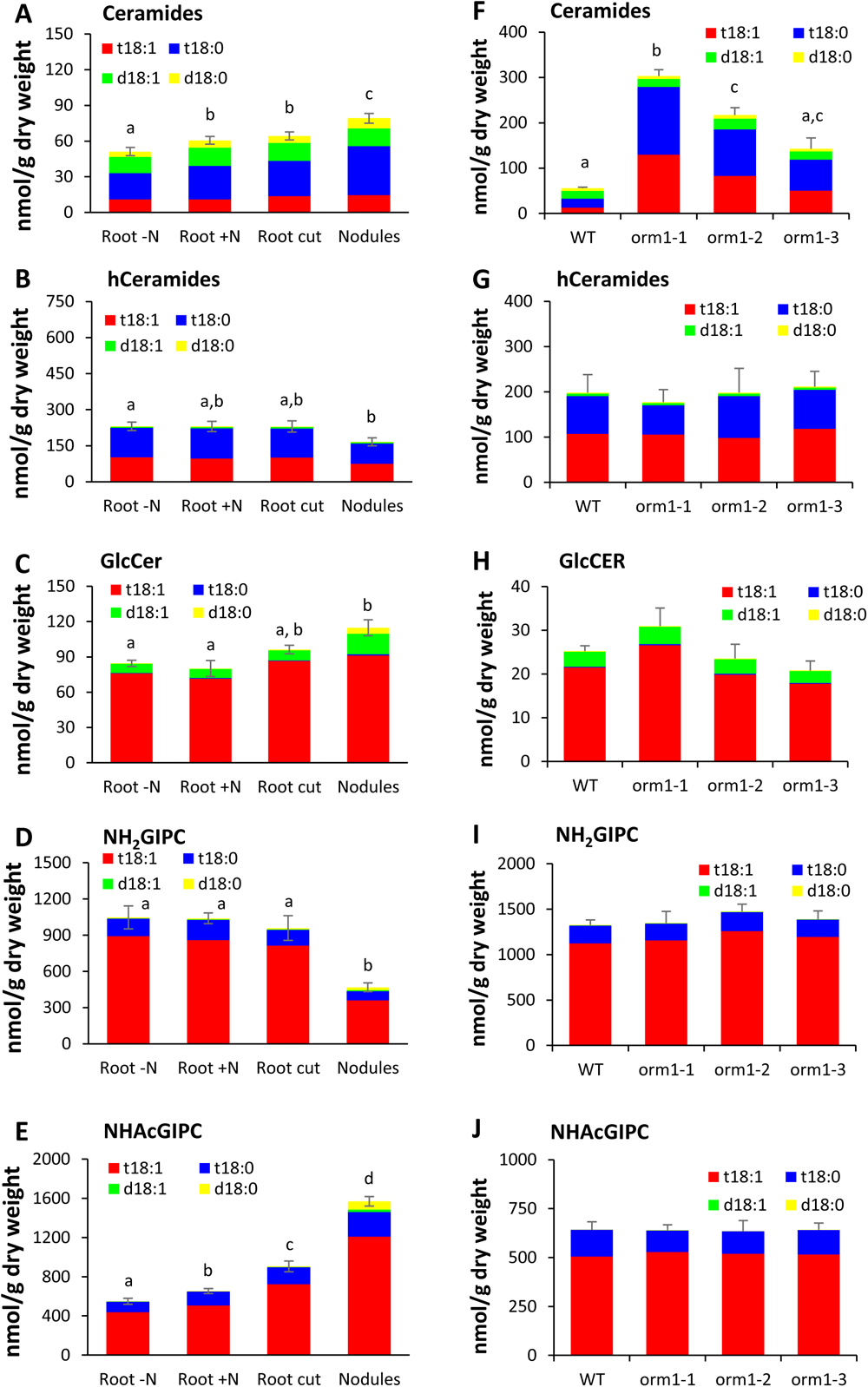
Mutations in *AeORM1* Cause Sphingolipid Accumulation in the Root. (**A**-**E**) Content of the following sphingolipid classes in roots and nodules of the WT plants after inoculation with *Bradyrhizobium* ORS278: (**A**) Ceramides, (**B**) hCeramides, (**C**) GlcCER, (**D**) NH2GIPC, (**E**) NHAcGIPC. Root-N: roots in BNM medium with 0.5 mM KNO3, Root+N: roots in BNM medium with 5 mM KNO3, Root cut: inoculated roots from which nodules were removed. (**F**-**J**) Content of the following sphingolipid classes in the roots of WT and *orm1* mutant plants: (**F**) Ceramides, (**G**) hCeramides, (**H**) GlcCer, (**I**) NH2GIPC, (**J**) NHAcGIPC. The LCB structures present in the analyzed shpingolipids were: d18:0, d18:1, t18:0 and t18:1. Values are means ± SE from three technical replicates. The experiments were performed twice with similar results. The result of one representative experiment are shown. Different letters represent significant differences determined using a Welch’s t-test, P < 0,05.

We pursued the sphingolipidomic profiling by determining sphingolipid composition in roots and nodules following inoculation with *Bradyrhizobium* strain ORS278 of WT plants. 4 weeks after inoculation, we excised the nodules from the roots and analyzed both organs. Sphingolipid amounts in uninoculated WT roots were simialr to inoculated WT roots from which the nodules had been excised, indicating that the presence of nitrogen-fixing nodules has no impact on the sphingolipid composition in the root (Figure 9A to E). Moderate differences were seen in the CER hCER and GlcCER composition between WT roots and nodules (Figure 9A-C). More striking were the detected opposed variations for the GIPC pool, with NH_2_GIPC (amino-GIPC) showing a 2 fold decrease and NHAcGIPC (acetylamino-GIPC) a 1.7 fold increase in WT nodules relative to WT roots (Figure 9D and E). As a result, NHAcGIPC constituted more that 75% of the total GIPC pool in WT nodules. Higher quantities of both NH_2_GIPC and NHAcGIPC d18:0-and d18:1-h16:0 were found in nodules. However, most of the variations actually concerned the predominant NH_2_GIPC and NHAcGIPC t18:0-and t18:1-h24:0 forms (Supplemental Figures 10 and 11).

Last, we compared the major classes of sphingolipids present in roots of the WT and three *orm1* mutant plants. The sphingolipid profiles of the *orm1* mutants showed an increased level in CER (*orm1-1*: ∼5 fold, *orm1-2*: ∼4 fold, *orm1-3*: ∼3 fold) as compared to the WT, which is specifically due to an overaccumulation of species containing trihydroxy LCB t18:0 and t18:1 (Figure 9F). Remarkably, the CER backbones containing very-long-chain fatty acids (VLCFAs) (C22 to C26, the C24 fatty acid being the major one) exhibited a much more dramatic increase than CER containing long-chain fatty acids (C16 and C20) (Supplemental Figure 6). No significant quantitative differences were observed for the other sphingolipid categories (Figure 9G-J). Overall, these findings indicated that the symbiotic and lateral root phenotypes of *orm1* mutants were associated with an overaccumulation of VLCFA-containing ceramids in *A. evenia* roots.

## Discussion

Using *A. evenia* mutants impaired in nodule formation, we identified the *AeORM1* gene that is predicted to encode an orosomucoid (ORM) protein. ORM proteins are key negative regulators of sphingolipids synthesis in eukaryotes and serve important functions in Arabidopsis, a nonsymbiotic plant, but a role forORM proteins in the rhizobial symbiosis has so far never been reported. This gene discovery illustrate how research on *A. evenia* nodulation is valuable to complement the information gained using historical model legumes.

In contrast to the recently identified *AeCRK* gene, which is essential for symbiosis establishment in *A. evenia* and that has no obvious ortholog in *M. truncatula* and *L. japonicus* (Quilbé et al., 2021), *ORM* genes are present in all legume species but in genetic screens in the historical model legumes thay have not been identified as important for nodulation. A reason for this could be that *A. evenia* constitutes a distinct genetic background. Indeed, while a single *ORM* gene copy in present in most legume species, including *M. truncatula* and *L. japonicus*, two paralogous genes are present in *A. evenia*. Differential retention of paralogs that results from the ancestral WGD event in Papilionoids has been repeatedly noticed in symbiotic genes. Notbale instances include AeSYMRK1&2, MtERN1&2, LjEIN1&2 (Quilbé et al., 2021; Yano et al., 2017; Miyata et al., 2013). Given the central role of ORM proteins in plant development, the inactivation in single copy-having legume species might be detrimental. In Arabidopsis, gene-edited *ORM* mutants yielded non-viable seeds, thereby impeding normal life cycle completion (Gonzales-Solis et al., 2020). In *A. evenia*, the presence of two *ORM* paralogs might provide functional redundancy or specialization. This idea is supported by the observation that *A. evenia orm1* mutants are still able to develop correctly and produce fertile seeds. Testing mutants for the single *ORM* gene in *M. truncatula* or *L. japonicus*, and for *AeORM2* -either alone or in combination with *AeORM1*-would help solving these questions. Another possibility is that *ORM* genes have differential symbiotic involvements in legumes. *A. evenia* symbiosis differs at several points from those present in *A. M. truncatula* and *L. japonicus*. It has a Nod factor-independent activation, and uses an intercellular infection and a Lateral Root Base (LRB) nodulation process (Quilbé et al., 2021, 2022b; Bonaldi et al., 2011). Interestingly, analysis of the *AeORM1* expression pattern revealed a constitutive expression at lateral root bases where rhizobial infection and nodule formation occur. This information complements the one recently obtained in the another LRB-nodulating legume, *A. hypogaea*, for which the expression of the symbiotic genes *AhNIN* and *AhCYCLOPS* was found to be induced specifically at lateral root bases upon rhizobial inoculation (Bhattacharjee et al., 2022). In addition, the early abortion of nodule development in the three *A. evenia orm1* mutants shows that *AeORM1* is essential for lateral root base-nodulation.

The *orm1* mutants produced bumps that differed from those previously described in certain *A. evenia* nodulation mutants for the *AePOLLUX*, *AeCYCLOPS* and *AeCRK* genes (Quilbé et al., 2022b) in being numerous and by having visible brown spots. Although a gradation in the severity of the nodulation phenotype was observed in the three *orm1* mutants, in all cases infection of plant cells by *Bradyrhizobium* clearly differed from WT plants both quantitatively (infected cells were less filled with bacteria) and qualitatively (the bacteria were not differentiated into bacteroids. Because nodule formation is triggered in the *orm1* mutants but rapidly arrested, we hypothesize that *AeORM1* is required for nodule development and not for early perception of bradyrhizobia or nodule inception. This conclusion is supported by a RT-qPCR analysis showing the partial induction of symbiotic genes following the inoculation of an *orm1* mutant with *Bradyrhizobium*, with the exception of the infection marker *AeVPY*. Interestingly, expression studies using a p*AeORM1*-GUS construct showed a specific expression of *AeORM1* in cells at the base of lateral roots where nodule primordia form following rhizobial infection. However, in nitrogen-fixing nodules, *AeORM1* expression was not associated with bacteroid-containing nodule cells. Based on these observations, we hypothesize that *AeORM1* is important for bacterial infection and nodule structure formation, acting downstream of nodule inception but prior to nodule differentiation. If right, *AeORM1* could represent a nodule emergence stage-specific regulator as described for *NF-YA1* (Nuclear Factor YA1) in *M. truncatula* and *L. japonicus* (Shrestha et al., 2020; Hossain et al., 2016; Laporte et al., 2014). The presence of brown spots in aborted nodules was found to correspond to the accumulation of phenolic compounds along with high concentrations of H_2_O_2_. They are reminiscent to those observed in mutants of different genes: *MtNAD1* (Nodules with Activated Defense 1), *MtDNF2* (Does Not Fix nitrogen 2), *MtSymCRK* (Symbiotic Cysteine-rich Receptor-loke Kinase), and *MtNIP/LATD* (Numerous Infections and Polyphenolics/Lateral roor-organ Defective) (Wang et al., 2016; Berrabah et al., 2014; Bourcy et al., 2013; Veereshlingam et al., 2004). Similar to these genes, *AeORM1* is likely to be important preventing inappropriate activation of defense-like responses during nodulation. Although aborted nodules with defense-like reactions was the most striking phenotype among *A. evenia orm1* mutants, playing closer attention also revealed an alteration in lateral root formation, all three mutants having lateral roots shorter as compared to WT plants. Such defect was not seen in other *A. evenia* nodulation mutants for *AeCCaMK* and *AeNSP2* (this study, Quilbé et al., 2022), comforting the idea that this phenotype is specific to mutations in *AeORM1*. In accordance, *AeORM1* is psecifically expressed in lateral root primordia and apex (prior and after emergence, respectively) in non-symbiotic conditions. Taken together, phenotypic and expression data for *AeORM1* establish a link between nodule and lateral root formation. Only a few genes shared between lateral root development and nodule formation are known, which include the aforementioned *MtNIP/LATD* gene that encodes a nitrate transporter (Bagchi et al., 2012; Yendrek et al., 2010) and *LBD16* (LOB-DOMAIN PROTEIN 16) coding for a transcription factor that acts with NF-YA1 to promote auxin signaling (Soyano et al., 2019; Schiessl et al., 2019). Although no role of ORM proteins in lateral root development has been described in non-legume plants, it is tempting to speculate that this developmental function has been co-opted for nodulation in legumes as for *MtNIP/LATD* and *LBD16*.

Given the very highl identity level of ORM proteins from legume and non-legume species, it is likey that ORM function is very conserved. ORM proteins are endoplasmic reticulum (ER)-resident membrane proteins that negatively regulate sphingolipid metabolism by forming a multiprotein complex with Serine PalmitoylTransferas (SPT). Tight regulation of sphingolipid homeostasis is critical since sphingolipids can act as bioactive molecules and are incorporated into cell membranes, influencing their structural and functional dynamics, especially vesicular trafficking, endocytosis and exocytosis. Therefore, they impact cell polarization, lipid raft formation, protein targeting to the membrane and also cell cytokinesis. We found that mutations in *AeORM1* primarily affected CER synthesis and composition in *A. evenia* roots leading to the accumulation of mainly VLCFAs-containing CER species (C22 to C26. In Arabidopsis, *ORM* gene inactivation also lead to the increase of ceramide content, but this concerns the ceramide backbones containing long-chain fatty acids (C16) rather than those with VLCFAs (Li et al., 2016; Kimberlin et al., 2016). These opposed data point to a differential impact of ORM mediation of SPT activity on ceramide synthase activities between *A. evenia* and Arabidopsis. These regulatory differences may be of importance in legume biology because VLCFAs and VLCFA-containing sphingolipids play a key role in cell proliferation, tissue patterning and lateral root development (Trinh et al., 2019; Roudier et al., 2010; Nagata et al., 2021). This suggests that a fine-tuning of VLCFA-Cer content in *A. evenia* may be essential for correct development of lateral roots and nodules. Furthermore, VLCFA-containing sphingolipids have been demonstrated to associate with Golgi-mediated protein-trafficking in Arabidopsis (Markham et al., 2011). Inactivation of ORM function may notably impact certain membrane-anchored proteins that intervene in the rhizobial symbiosis, such as the Golgi-located VPY (Liu et al., 2019). Recent work in *M. truncatula* revealed that sphingolipids are also important for bacterial accommodation as they are part of the plant-produced membrane delimiting symbiosomes containing the bacteria. Bacterial accommodation is linked to the reprogramming of sphingolipid glycosylation (Moore et al., 2021). These changes were shown to be mediated by *MtGINT1* (Glucosamine Inositol Phosphorylceramide Transferase 1) whose inactivation impaired both nodulation and mycorrhization. In *A. evenia*, a similar change in sphingolipid glycosylation between roots and nodules was shown here, indicating that this process may be a general rule in legumes. In contrast to *M. truncatula gint1* silenced lines, *A. evenia orm1* mutants were not altered for the mycorrhizal symbiosis. Determining the precise cellular function of ORM-regulated sphingolipid homeostasy in nodule formation and how it relates to lateral root formation represents an important avenue in the research to uncover the mechanisms of intercellular infection and Lateral Root Base nodulation in *A. evenia* and, the rhizobial symbiosis in legumes in general.

## Materials and methods

### Plant materials and growth conditions

In this study, *Aeschynomene evenia* CIAT22838 was used as wild-type reference plant and in some experiments its *ccamk*-3 mutant as negative control (Quilbé et al., 2021, 2022b). All analyzed *orm* mutants were isolated from the EMS-mutagenized collection of *Aeschynomene evenia* CIAT22838 plants that have been shown to have phenotypic defects in nodulation (Quilbé et al., 2021). *A. evenia* seeds were scarified for 40 min with sulfuric acid (96%) and rinsed several times with distilled water; whereafter germination was induced by overnight incubation in distilled water containing 0.01% (v/v) ethrel (BAYER). For root phenotyping and seed production, 1-day-old seedlings were transferred to plastic pots filled with attapulgite whereafter the plants were cultivated in the greenhouse (28°C and 70% relative humidity) as detailled in Quilbé et al. (2021).

### Analysis of the root system architecture

Plants were cultured in tubes filled with BNM medium as published (Quilbé et al., 2022b). Roots were scanned using an EPSON GT-15000 scanner and measurements of root length or density were performed using the Optimas 6.1 software (Media Cybernitics, Silverspring, MD, USA).

### Plant nodulation and acetylene reduction assays

For analysis of rhizobial infection and nodulation, *Bradyrhizobium* WT strain ORS278 and the derivative strains ORS278-GUS or ORS278-GFP were used (Bonaldi et al., 2011; Giraud et al., 2007). In each case, nodulation tests were performed in covered tubes to protect roots from light. Bacterial culture, root inoculation, nitrogenase enzyme activity (through the measurement of acetylene reducing activity - ARA) and macroscopic observations were performed as already published (Quilbé et al., 2022b; Arrighi et al., 2012; Bonaldi et al., 2011).

For confocal microscopy, root sections containing nodules from plants inoculated with ORS278-GFP were harvested and rinsed with distilled water. The rinsed root sections were embedded in 5% agar, and sectioned (70 µM) using a vibratome (VT1000S; Leica, Nanterre, France). Nodule sections induced by ORS278-GFP were analyzed as described by Bonaldi et al. (2011).

### Plant mycorrhization

Mycorrhization tests were performed by inoculating 5 day-old *A. evenia* seedlings with *Rhizophagus irregularis* DAOM197198 (Agronutrition, Carbonne, France) and by optimizing culture conditions described in Quilbé et al. (2022b). In short, plants were watered three times a week with the nutritive solution, trays containing pots with plants were changed weekly and plants were cultured for 6 weeks following inoculation with spores. Fungal colonization was assessed on six plants/genotype/biological repeat using the Myco-Calc method as detailed in Quilbé et al. (2022b).

### Histochemical staining

Bacterial infection of plant tissue and nodule development were analyzed using 70 µm-thick vibratome (Leica VT1000S) sections of freshly harvested material. In case of inoculation with strain ORS278-GUS, sections were stained with X-Gluc (Fabre et al., 2015). Root sections were observed using a Nikon DS-Ri2 microscope either under bright field illumination or using the GFP and mCherry filters to analyze autofluorescence. To estimate H_2_O_2_ production *in situ*, root and nodule sections were immersed in a 1 mg.ml^-1^ of 3’,3’-diaminobenzidine (DAB) solution, vacuum infiltrated for 2 min and then incubated for 2-3h at 25°C before observation. The accumulation of phenolic compounds was detected by potassium permanganate-methylene blue staining as described (Bourcy et al., 2013).

### Genetic characterization and sequencing of *A. evenia* mutants

For the genetic determinism analysis of nodulation mutants, plants were manually hybridized with the wild type line. Approximately 600 plants of the mutant x WT F2 progeny were cultured in the greenhouse, inoculated with the *Bradyrhizobium* strain ORS278 and roots were phenotyped 4 weeks post inoculation. For each nodulation mutant, DNA was extracted from pooled roots of 95 to 150 F2 plants with a mutant phenotype using the CTAB method. Library preparation and sequencing on a NovaSeq sequencer was performed at the Norwegian Sequencing Center (CEES, Oslo, Norway). The 150 bp-paired end reads were processed to conduct the Mapping-by-Sequencing approach to identify candidate genes as previously described (Quilbé et al., 2021). Allelism tests were performed by directed crossing of the nodulation mutants and root phenotyping of the F1 progeny cultured either in the greenhouse or growth chamber and inoculated with the *Bradyrhizobium* strain ORS278. Genetic characteristics of the mutants are provided in Supplemental Tables 1 and 2.

### *In silico* gene analysis

Genes homologous to *AeORM1* were identified in legume species by mining the orthogroups database generated with OrthoFinder (Quilbé et al., 2021). The dataset was completed by searching for other legume genes as well as Arabidopsis and rice *ORM* genes in the Legume Information System (https://www.legumeinfo.org), The Arabidopsis Information Resource (https://www.arabidopsis.org) and the Rice Genome Annotation Project (http://rice.uga.edu) databases, respectively. A ML phylogenetic tree reconstruction was obtained by aligning nucleotide sequences with the MUSCLE program that is incorporated in the MEGA X (v10.1.8) software. Aligned sequences were further processed in MEGA X using the maximum likelihood approach and the Kimura 2-parameter model with a 1000x bootstrap (BS). The data are presented as rooted tree using rice as outgroup. For the *AeORM1* and *AeORM2* genes, microsynteny analysis was performed using the Legume Information System with the Genome Context Viewer (https://legumeinfo.org/lis_context_viewer) to visualize the gene collinearity in syntenic regions and their expression patterns were obtained using the *A. evenia* gene atlas available at the AeschynomeneBase (http://aeschynomenebase.fr/content/gene-expression).

Protein sequences were aligned using the MUSCLE program in the MEGA X software and sequence alignments were visualized with Jalview v2.11.0. AeORM1 protein domains were identified, annotated using InterProscan (http://www.ebi.ac.uk/interpro/) and DeepTMHMM (https://dtu.biolib.com/DeepTMHMM), and refined by comparative structural analysis with other *Arabidopsis* ORM proteins. Amino acid conservation and properties were further analyzed with Weblogo3 (https://weblogo.threeplusone.com).

### Constructs for *in planta* gene analyses

For the analysis of the *AeORM1* gene, different constructs were generated based on the Golden Gate cloning method using vectors and modules generated by Fliegmann et al. (2016). In a first step, sequences of the *AeORM1* 2.166-bp native promoter (pro*AeORM1*) and of the full-length CDS of *AeORM1* containing either the stop codon or not, were flanked by two *Bsa*I sites designated so as to generate specific cohesive 5’ protuding ends. These sequences were synthetized and cloned into Puc57-*Bsa*I-free plasmid by GeneCust (www.genecust.com) and checked by DNA sequencing. For *A. evenia* root transformation, the cloned promoter pro*AeORM1* was fused to the *GUS* gene and the full-length CDS *AeORM1* with stop codon by GoldenGate cloning, using a vector based on pCambia2200 expressing DsRED.

### Functional complementation experiment

Using *Agrobacterium rhizogenes*-mediated transformation (Quilbé et al., 2021), seedlings of *orm1-1* were transformed using ARqua1 strains containing either empty vector (EV) or ProAeORM1:AeORM1 construct in pCambia2200DsRED. Plants were initially cultured on agar plates with half-strength MS medium (Murashige and Skoog basal salt mixture). After three weeks of growth, transformed roots were identified by expression of the DsRed marker and non-transformed ones were removed from the plants. Subsequently, plants were transfered to Falcon tubes filled with BNM liquid medium and inoculated with the *Bradyrhizobium* strain ORS278. Nodulation was quantified 28 days after inoculation. The number of nodulated plants out of the total transformed plants and the number of nodules per nodulated plant were determined.

### Analysis of promoter-GUS expressing plants

The pCambia2200DsRED vector containing the proAeORM1-GUS fusion was transformed into *Agrobacterium rhizogenes* strain ARqua1, and cells harboring this plasmid were used to transform WT *A. evenia* as described (Quilbé et al., 2021). Plants developing transformed roots were selected as described above. The *AeORM1* promoter activity was analyzed in non-inoculated roots and at different times after inoculation with the *Bradyrhizobium* strain ORS278. Untransformed WT seedlings were used as negative control. Whole roots and 70-µm-thick nodule sections (obtained with a Leica VT1000S vibratome (Nanterre, France) were stained with X-gluc (Fabre et al., 2015). Images of GUS-analyzed tissues were taken with a stereo-macroscope (Niko AZ100, Champigny-sur-Marne, France) using the Nikon Advanced software.

### RNA isolation and quantitative RT-PCR

For expression analysis of *AeORM1* and *AeORM2*, RNA material that have been previously generated in biological triplicates for the WT line in non-inoculated conditions or inoculated either with *Bradyrhizobium* strain ORS278 or *Rhizophagus irregularis* DAOM197198 was used (Quilbé et al., 2022b). For expression analysis of symbiotically induced genes, pools of 5 roots were collected per genotype and time-point to serve as source of RNA material. They were obtained across four independent experiments. Total RNA extraction from roots, reverse transcription quantitative PCR (qRT-PCR) were performed as described (Gully et al., 2018; Fabre et al., 2015). Expression levels were normalized with the *AeEF1-*α and *AeUbi* reference genes. Gene-specific primers were designed with Beacon Designer (Premier Biosoft) (Supplemental Table 3).

### Sphingolipid extraction and analysis

For root sphingolipid analysis, roots from 4-week-old plants grown in liquid BNM medium (supplemented with 0.5 or 5 mmol KNO_3_) were harvested and three pools of three roots were formed. For nodule sphingolipid analysis, nodulated roots of plants inoculated with *Bradyrhizobium* ORS278 were collected at 4wpi and nodules were separated from 10 roots to constitute three bulks of roots and nodules, respectively. The experiments were performed twice. Sphingolipids were extracted from 6-10 mg of lyophilized plant material as described previously (Tellier et al., 2014). Here, 1 mL of extraction solvent (isopropanol:hexane:water, 55:20:25) and 10 μL of adequate internal standards were added to 2 mg of freeze-dried material and grinded using a Polytron homogenizer. The sample was incubated at 60°C for 15 min. After centrifugation at 1620 *g* for 5 min, the supernatant was recovered and the pellet extracted once more with 1 mL of extraction solvent as previously. Supernatants were combined and dried using a Speed-Vac evaporator. To improve ionisation, the samples were subjected to alkaline hydrolysis whereafter, they were re-suspended in 100 μL of tetrahydrofuran (THF):methanol:water (2:1:2) containing 0.1% formic acid (for GIPCs classes and GlcCers) or THF (for Cers and hCers) by sonication and filtrated before analysis. Sphingolipid standards used were GM1 (20 nmol), C12-GlcCer (10 nmol) and C12-Cer (1 nmol) in 1 mL of extraction solvent (isopropanol:hexane:water, 55:20:25). They were purchased from AvantiPolar Lipids Inc. (Alabaster, AL, USA). Ultra high-performance liquid chromatography (UPLC)-electrospray ionisation (ESI)-tandem mass spectrometry (MS/MS) analyses were carried out on a Waters Acquity UPLC system coupled to a Waters Xevo tandem quadrupole mass spectrometer (Manchester, UK) equipped with an ESI source. The mass analyses were performed in the positive multiple reaction monitoring (MRM) mode. Chromatographic conditions and mass spectrometric parameters were defined previously (Tellier et al., 2014).

### Statistical analyses

To evaluate statistically the monogenic determinism of the three *orm1* mutants a Student’s *t*-test was performed t. A Mann-Whitney test was used to compare gene expression levels in the WT and *orm1-1* lines. One-way Kruskal-Wallis analysis was performed to compare root parameters between the WT line and the three *orm1* mutants. Values of P < 0.05 were considered statistically significant. A Welch’s *t*-test was used to compare sphingolipid levels between *orm1* mutants and the WT line. The statistical analyses were done using the R package or GraphPad Prism 8.3.0.

### Accession numbers

The Mapping-by-Sequencing data generated for the *orm1* mutant in this study were deposited in the NCBI database under BioProject ID: PRJNA727694.

## Supplemental Data

**Supplemental Figure S1.** Identification of *orm1* mutant alleles by Mapping-by-Sequencing.

**Supplemental Figure S2.** Syntenic localization of *ORM* genes in *A. evenia*.

**Supplemental Figure S3.** Structure and alignment of plant ORM proteins.

**Supplemental Figure S4.** *AeORM1* and *AeORM2* are expressed throughout the plant.

**Supplemental Figure S5.** Nodulation kinetics of the WT and *orm1* mutant plants after inoculation with *Bradyrhizobium* ORS278.

**Supplemental Figure S6.** Comparison of NHAcGIPC and NH_2_GIPC molecular species composition between the WT roots and nodules after inoculation with Bradyrhizobium ORS278.

**Supplemental Figure S7.** Ceramide molecular species composition representing the exact pairings of LCB and fatty acid in roots of WT and *orm1* mutant plants.

**Supplemental Table S1.** Genetic data of the *Aeschynomene evenia orm1* mutants.

**Supplemental Table S2.** Allelism analysis of the *Aeschynomene evenia orm1* mutants.

**Supplemental Table S3.** Complementation of the nodulation phenotype of *orm1-1* mutant plants by expressing WT *AeORM1*.

**Supplemental Table S4.** Primer sequences used for qRT-PCR.

## Acknowledgements

We thank Robin Duponnois (LSTM Laboratory, IRD) for assistance with the characterization of *A. evenia* nodulation mutants and for kindly providing fungi spores for mycorrhization experiments. We also thank Benoit Lefebvre (LIPME Laboratory, INRAE) for providing the pCambiaRedGG plasmid for GoldenGate cloning.

## Author contributions

NN, FG, and JFA conceived and designed the experiments; NN, MP, FEM, FT, MR, NHA, CK, FG, and JFA performed the experiments and analyzed the data; and NN, FG, and JFA. wrote the article.

The author responsible for distribution of materials integral to the findings presented in this article in accordance with the policy described in the Instructions for Authors (https://academic.oup.com/plphys) is: Jean-François Arrighi.

## References

1. Alsiyabi A, Solis AG, Cahoo E.B, Saha R. (2021) Dissecting the regulatory roles of ORM proteins in the sphingolipid pathway of plants. PLoS Comput Biol. 17:e1008284

2. Arrighi JF, et al. (2012) *Aeschynomene evenia*, a model plant for studying the molecular genetics of the nod-independent rhizobium-legume symbiosis. Molecular Plant Microbe Interactions. 25: 851–61

3. Bagchi R, Salehin M, Adeyemo OS, Salazar C, Shulaev V, Sherrier DJ, Dickstein R. (2012) Functional assessment of the Medicago truncatula NIP/LATD protein demonstrates that it is a high-affinity nitrate transporter. Plant Physiol. 160:906–16

4. Bhattacharjee O, Raul B, Ghosh A, Bhardwaj A, Bandyopadhyay K, Sinharoy S. (2022) Nodule INception-independent epidermal events lead to bacterial entry during nodule development in peanut (*Arachis hypogaea*). New Phytol. 236:2265–2281

5. Berrabah F, et al. (2019) Insight into the control of nodule immunity and senescence during *Medicago truncatula* symbiosis. Plant Physiol. 191: 729–746

6. Berrabah F, et al. (2014) A nonRD receptor-like kinase prevents nodule early senescence and defense-like reactions during symbiosis. New Phytol. 203:1305–1314

7. Bonaldi K, Gargani D, Prin Y, Fardoux J, Gully D, Nouwen N, Goormachtig S, Giraud, E. (2011) Nodulation of *Aeschynomene afraspera* and *A. indica* by photosynthetic *Bradyrhizobium* Sp. strain ORS285: the nod-dependent versus the nod-independent symbiotic interaction. Molecular Plant Microbe Interactions 24: 1359–71

8. Bourcy M, Brocard L, Pislariu CI, Cosson V, Mergaert P, Tadege M, Mysore K.S, Udvard M.K, Gourion B, Ratet P. (2013) *Medicago truncatula* DNF2 is a PI-PLC-XD-containing protein required for bacteroid persistence and prevention of nodule early senescence and defense-like reactions. New Phytol. 197: 1250–1261

9. Capoen W, Oldroyd G, Goormachtig S, Holsters M. (2010). *Sesbania rostrata*: a case study of natural variation in legume nodulation. New Phytol 186: 340–5

10. Chaintreuil C, et al. (2018) Naturally occurring variations in the nod-independent model legume *Aeschynomene evenia* and relatives: a resource for nodulation genetics. BMC Plant Biol. 18: 54

11. Fabre S, Gully D, Poitout A, Patrel D, Arrigh J.F, Giraud E, Czernic P, Cartieaux F. (2015) Nod factor-independent nodulation in *Aeschynomene evenia* required the common plant microbe symbiotic toolkit. Plant Physiol. 169: 2654L2664

12. Feng J, Lee T, Schiessl K, Oldroyd GED. (2021) Processing of NODULE INCEPTION controls the transition to nitrogen fixation in root nodules. Science 374: 629–632

13. Fliegmann J, Jauneau A, Pichereaux C, Rosenberg C, Gasciolli V, Timmers AC, Burlet-Schiltz O, Cullimore J, Bono JJ. (2016) LYR3, a high-affinity LCO-binding protein of Medicago truncatula, interacts with LYK3, a key symbiotic receptor. FEBS Lett. 590: 1477–87

14. Giraud E, et al. (2007) Legumes symbioses: absence of *Nod* genes in photosynthetic bradyrhizobia. Science 316: 1307–1312

15. Gobbato E. (2015) Recent developments in arbuscular mycorrhizal signaling. Cur Op Plant Biol 26: 1–7

16. Gonzalez-Solis A, Han G, Gan L, L Y, Markham, JE, Cahoon E, Dunn T.M, Cahoon EB. (2020) Unregulated sphingolipid biosynthesis in gene-edited Arabidopsis ORM mutants results in nonviable seeds with strongly reduced oil content. Plant Cell. 32:2474–2490

17. Gully D, et al. (2018) Transcriptome profiles of Nod factor-independent symbiosis in the tropical legume *Aeschynomene evenia*. Sci Rep 8: 10934

18. Hossain MS, et al. (2016) *Lotus japonicus* NF-YA1 plays an essential role during nodule differentiation and targets members of the SHI/STY gene family. Mol Plant Microbe Interact. 29:950–964

19. Kimberlin AN, Han G, Luttgeharm KD, Chen M, Cahoon RE, Stone JM, Markham J., Dunn TM, Cahoon EB. (2016) ORM expression alters sphingolipid homeostasis and differentially affects ceramide synthase activity. Plant Physiol. 172: 889–900

20. Lace B, Su C, Invernot Perez D, Rodriguez-Franco M, Vernié T, Batzenschlager M, Egli S, Liu CW, Ott T. (2023) RPG acts as a central determinant for infectosome formation and cellular polarization during intracellular rhizobial infections. Elife. 12: e80741

21. Laporte P, et al. (2014) The CCAAT box-binding transcription factor NF-YA1 controls rhizobial infection. J Exp Bot. 65:481–94

22. Li J, Yin J, Rong C, Li KE, Wu JX, Huang LQ, Zeng HY, Sahu SK, Yao N. (2016) Orosomucoid proteins interact with the small subunit of serine palmitoyltransferase and contribute to sphingolipid homeostasis and stress responses in Arabidopsis. Plant Cell. 28: 3038–3051

23. Li QG, Zhang L, Li C, Dunwell JM, Zhan Y.M. (2013) Comparative genomics suggests that an ancestral polyploidy event leads to enhanced root nodule symbiosis in the Papilionoideae. Mol Biol Evol. 30: 2602–2611

24. Liu CW, et al. (2019) A protein complex required for polar growth of rhizobial infection threads. Nat Commun. 10: 2848

25. Markham JE, Molino D, Gissot L, Bellec Y, Hématy K, Marion J, Belcram K, Palauqui JC, Satiat-Jeunemaître B, Faure JD. (2011) Sphingolipids containing very-long-chain fatty acids define a secretory pathway for specific polar plasma membrane protein targeting in Arabidopsis. Plant Cell. 23:2362–78

26. Miyata K, Kawaguchi,M, Nakagawa T. (2013) Two distinct EIN2 genes cooperatively regulate ethylene signaling in *Lotus japonicus*. Plant Cell Physiol. 54:1469–77

27. Moore W, Chan C, Ishikawa T, Rennie EA, Wipf HM, Benites V, Kawai-Yamada M, Mortimer JC, Scheller HV. (2021) Reprogramming sphingolipid glycosylation is required for endosymbiont persistence in *Medicago truncatula*. Curr Biol. 31: 2374–2385.e4

28. Nagata K, Ishikawa T, Kawai-Yamada M, Takahashi T, Abe M. (2021) Ceramides mediate positional signals in *Arabidopsis thaliana* protoderm differentiation. Development. 148:dev194969

29. Quilbé J, Montiel J, Arrighi JF, Stougaard J. (2022a) Molecular mechanisms of intercellular rhizobial infection: novel findings of an ancient process. Front Plant Sci. 13: 922982

30. Quilbé J, Nouwen N, Pervent M, Guyonnet R, Cullimore J, Gressent F, Araújo NH, Gully D, Klopp C, Giraud E, Arrighi JF. (2022b) A mutant-based analysis of the establishment of Nod-independent symbiosis in the legume *Aeschynomene evenia*. Plant Physiol. 190: 1400–1417

31. Quilbé J, et al. (2021) Genetics of nodulation in *Aeschynomene evenia* uncovers mechanisms of the rhizobium-legume symbiosis. Nat Communications. 12: 829

32. Roy S, Liu W, Nandety RS, Crook ., Mysore KS, Pislariu CI, Frugoli ., Dickstein R, Udvardi MK. (2020) Celebrating 20 years of genetic discoveries in legume nodulation and symbiotic nitrogen fixation. Plant Cell. 32: 15–41

33. Roudier F, et al. (2010) Very-long-chain fatty acids are involved in polar auxin transport and developmental patterning in Arabidopsis. Plant Cell. 22:364–75

34. Sharma V, Bhattacharyya S, Kumar R, Kumar A, Ibañez F, Wang J, Guo B, Sudini HK, Gopalakrishnan S, DasGupta M. (2020) Molecular basis of root Nodule symbiosis between *Bradyrhizobium* and ‘Crack-Entry’ legume groundnut (*Arachis hypogaea* L.). Plants (Basel). 9: 276

35. Schiessl K, et al. (2019) NODULE INCEPTION recruits the lateral root developmental program for symbiotic nodule organogenesis in *Medicago truncatula*. Curr Biol. 29: 3657–3668.e5

36. Shrestha A, et al. (2021) *Lotus japonicus* Nuclear Factor YA1, a nodule emergence stage-specific regulator of auxin signalling. New Phytol. 229:1535–1552

37. Soyano T, Shimoda Y, Kawaguchi M, Hayashi M. (2019) A shared gene drives lateral root development and root nodule symbiosis pathways in Lotus. Science. 366: 1021–1023

38. Tellier F, Maia-Grondard A, Schmitz-Afonso I, Faure JD. (2014) Comparative plant sphingolipidomic reveals specific lipids in seeds and oil. Phytochemistry. 103: 50–58

39. Teulet A, et al. (2019) The rhizobial type III effector ErnA confers the ability to form nodules in legumes. Proc Natl Acad Sci U S A. 116: 21758–21768

40. Trinh DC, et al. (2019) PUCHI regulates very long chain fatty acid biosynthesis during lateral root and callus formation. Proc Natl Acad Sci U S A. 116:14325–14330

41. Veereshlingam H, Haynes JG, Penmetsa RV, Cook DR, Sherrier DJ, Dickstein R. (2004) *Nip*, a symbiotic *Medicago truncatula* mutant that forms root nodules with aberrant infection threads and plant defense-like response. Plant Physiol. 136:3692–702

42. Wang C, et al. (2016) NODULES WITH ACTIVATED DEFENSE 1 is required for maintenance of rhizobial endosymbiosis in *Medicago truncatula*. New Phytol. 212:176–91

43. Yano K, Aoki S, Liu M, Umehara Y, Suganuma N, Iwasaki W, Sato S, Soyano T Kouchi H, Kawaguchi M. (2017) Function and evolution of a *Lotus japonicus* AP2/ERF family transcription factor that is required for development of infection threads. DNA Res. 24:193–203

