## Supplemental Figures 1-7 for "*ORM*-mediated regulation of sphingolipid biosynthesis is essential for nodule formation in *Aeschynomene evenia*"

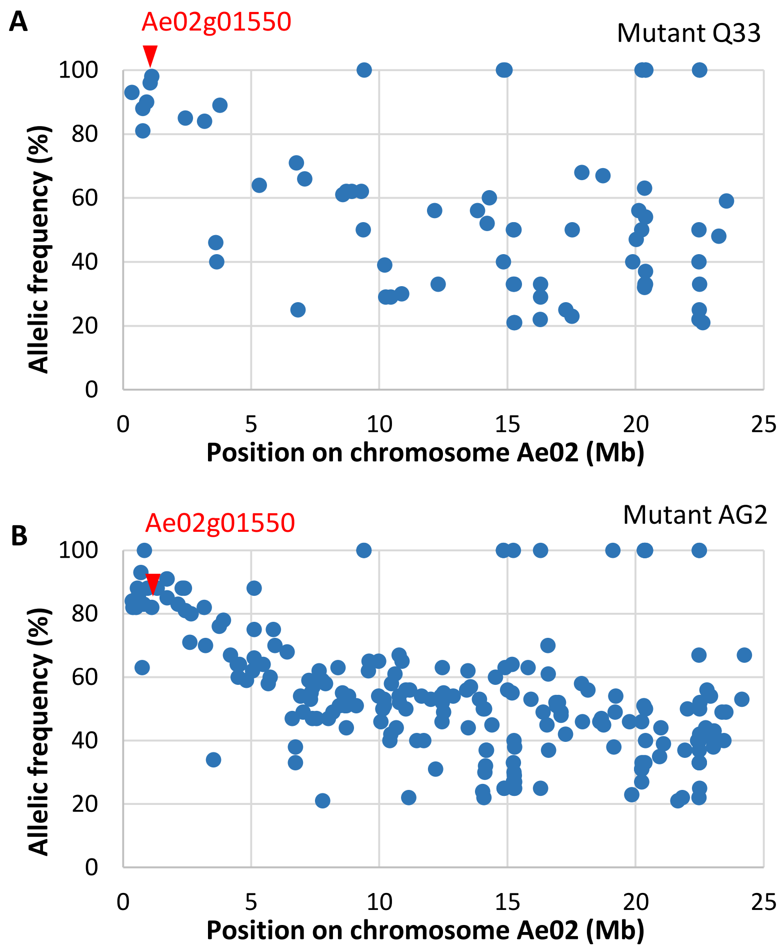


**Supplemental Figure S1.** Identification of *orm1* mutant alleles by Mapping-by-Sequencing. Frequency of EMS-induced mutations in bulks of *A. evenia* mutant backcrossed F2 plants determined with the Mapping-by-Sequencing approach: (**A**) Q33 mutant and (**B**) AG2 mutant. A genetic linkage corresponding to a shift in the allelic frequency is visible at the beginning of chromosome Ae02. The SNPs identified in the Ae02g01550 gene (*AeORM1*) are indicated by red arrows (AF= 98% and 82% for the Q33 and AG2 mutants, respectively) and are expected to be responsible for the observed symbiotic phenotype.

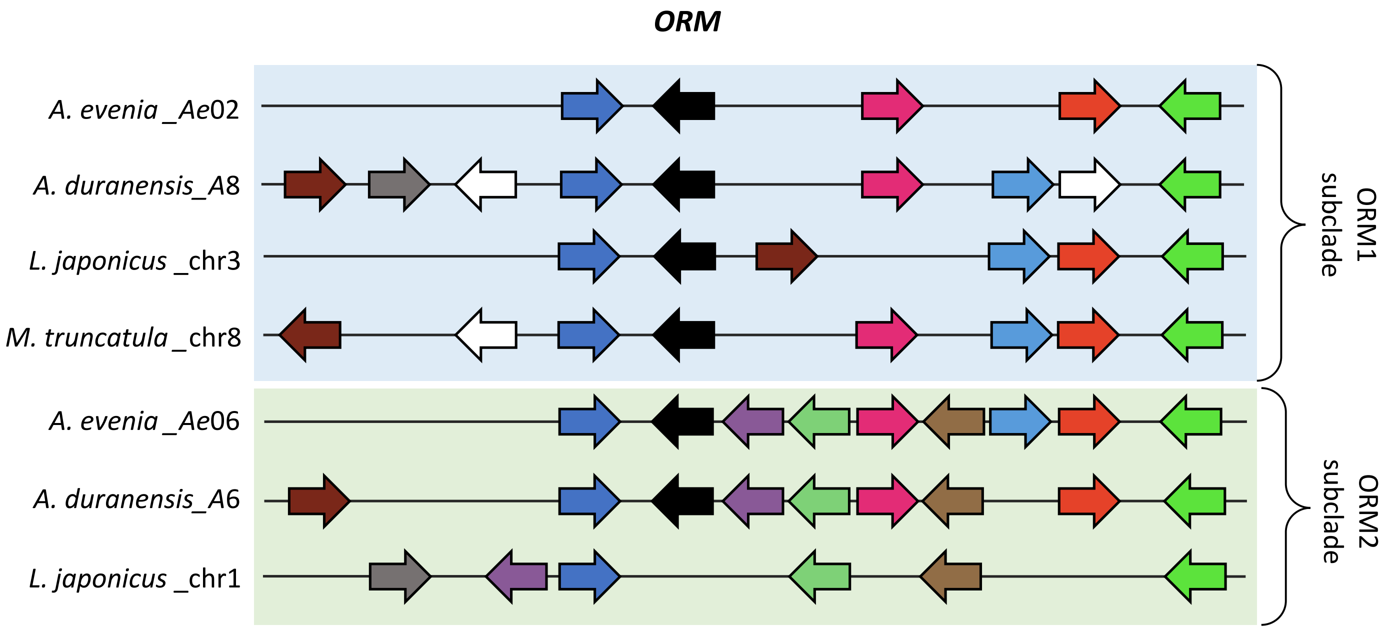


**Supplemental Figure S2.** Syntenic localization of *ORM* genes in *A. evenia*. Schematic representation of the microsynteny analysis of *ORM* genes (black arrow) present in *A. evenia*, *A. duranensis, L. japonicus* and *M. truncatula*. Orthologous/paralogous gene pairs are indicated through the use of a common colour. For clarity, orphan genes are not indicated in the figure. Green and blue rectangles highlight the duplicated regions putatively derived from the ~58 MYA WGD event in Papilionoid legumes. Note the absence of an *ORM* gene of the ORM2 subclade in *L. japonicus* and *M. truncatula* but the retention of the syntenic region in *L. japonicus*.

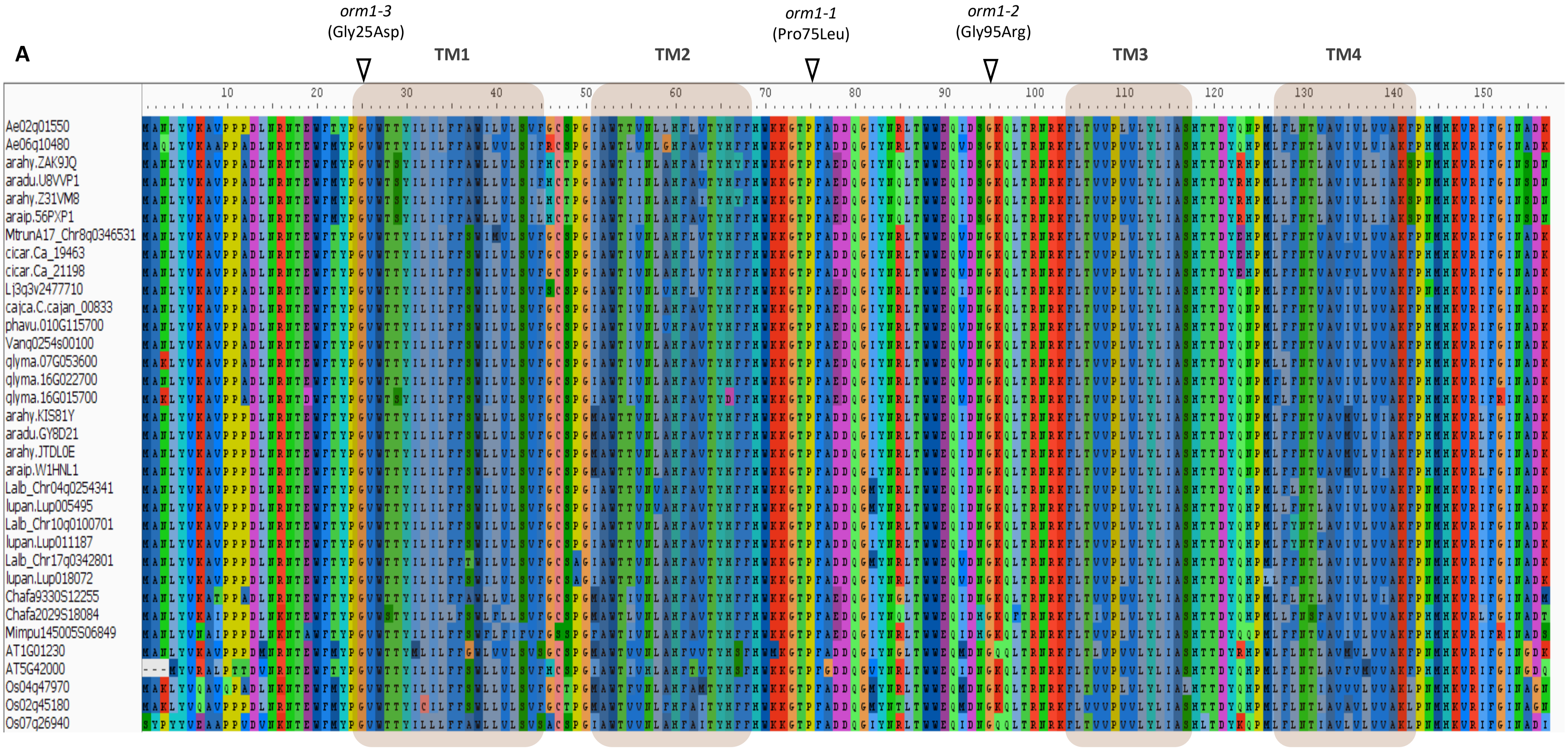


**Supplemental Figure S3.** Structure and alignment of plant ORM proteins. (Supports Figure 2) MUSCLE protein alignment of ORM proteins from selected species: *Aeschynomene evenia* (Ae), *Arabidopsis thaliana* (AT), *Arachis hypogaea* (Arahy), *Arachis duranensis* (Aradu), *Arachis ipaiensis* (Araip), *Cajanus cajan* (C. cajan), *Chamaecrista fasciculata* (Chafa), *Cicer arietum* (Ca), *Glycine max* (Glyma), *Lupinus albus* (Lalb), *Lupinus angustifolius* (Lup), *Lotus japonicus* (Lj), *Medicago truncatula* (Mtrun), *Mimosa pudica* (Mimpu), *Oryza sativa* (Os), *Phaseolus vulgaris* (Phvul), *Vigna angularis* (Vang). Predicted transmembranes (TM) are indicated by grey boxes and the position of AA changes resulting from EMS mutations in *AeORM1* are indicated by arrow heads.

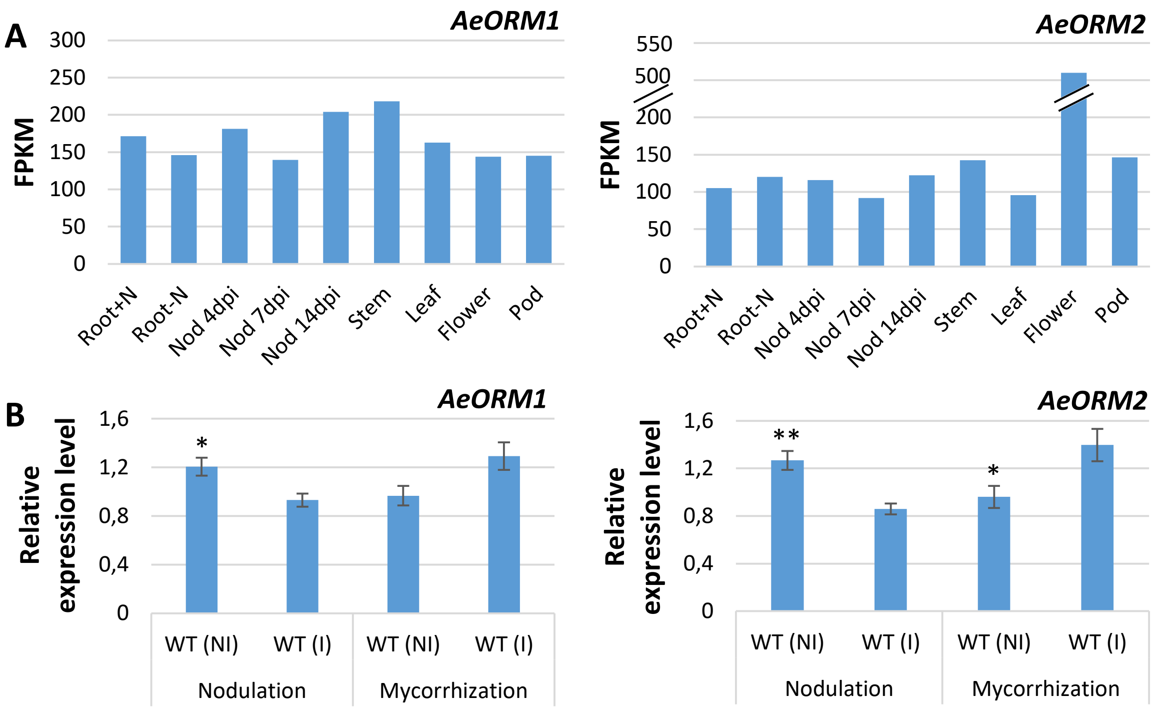


**Supplemental Figure S4.** *AeORM1* and *AeORM2* are expressed throughout the plant. (**A**) *AeORM1* and *AeORM2* expression in different tissues of *A. evenia* plants as found in the *A. evenia* Gene Atlas. Expression is given in normalized FPKM read counts. (**B**) Relative expression levels of *AeORM1* and *AeORM2* during nodulation and mycorrhization. Expression levels were determined in non-inoculated (NI) or inoculated (I) WT plants by RT-qPCR analysis. For nodulation, expression was determined at 7 days post-inoculation with *Bradyrhizobium* ORS278 and for mycorrhization at 8 weeks post-inoculation with *R. irregularis*. Expression values were normalized using *AeEF1a* and *Ubiquitin* expression levels as standard. Means and standard deviation (s.d.) were derived from three to four biological replicates. Asterisks indicate significant differences (*P < 0.05, **P < 0.01 ; Student's t-test) between the two conditions.

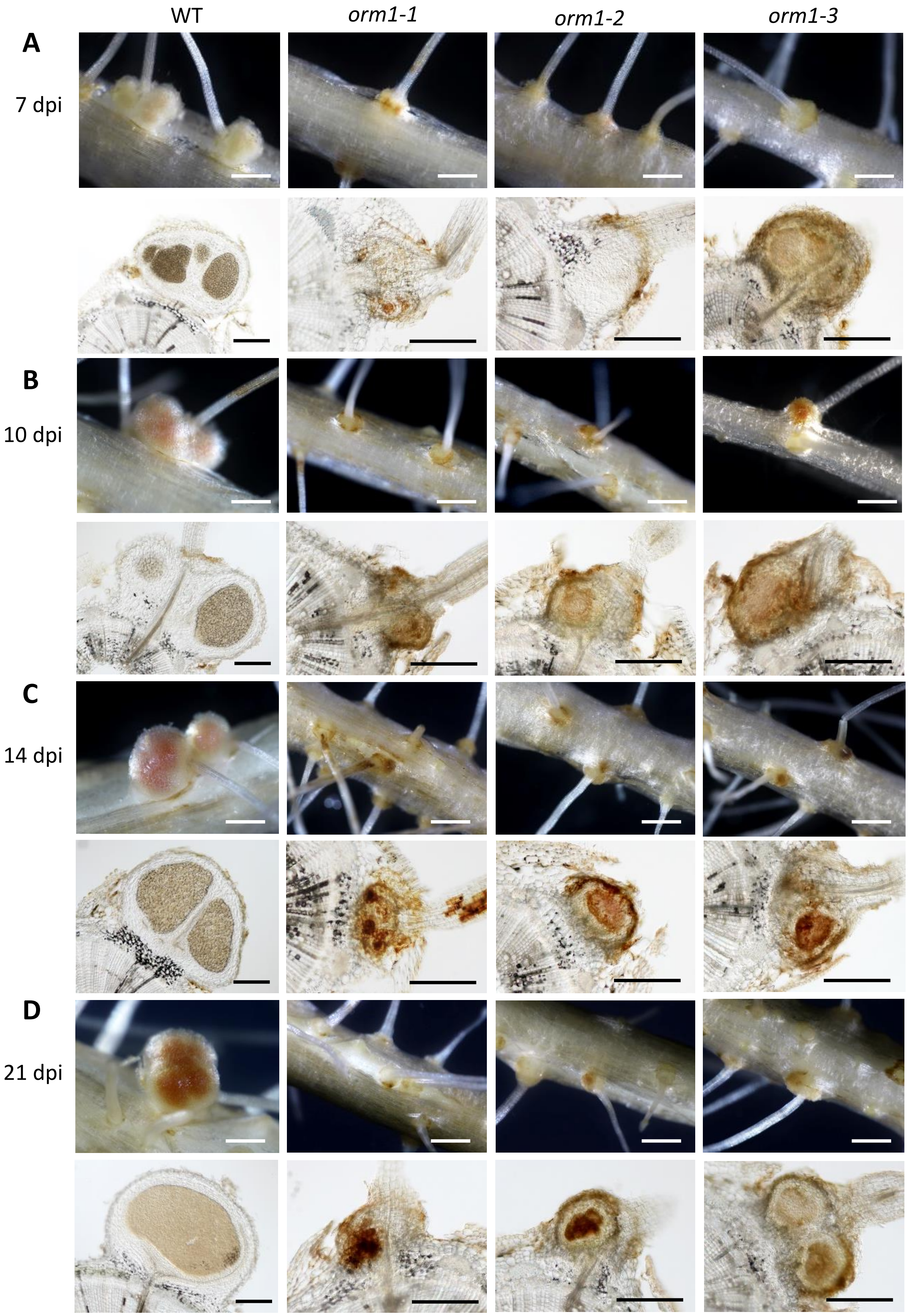


**Supplemental Figure S5.** Nodulation kinetics of the WT and *orm1* mutant plants after inoculation with *Bradyrhizobium* ORS278. Whole roots and 70 µm-thick root or nodule sections (upper and lower panels, respectively) of WT and *orm1* mutant plants at (**A**) 7 dpi, (**B**) 10 dpi, (**C**) 14 dpi, and (**D**) 21 dpi with *Bradyrhizobium* ORS278. Arrows show bumps and/or brown spots at nodulation sites. Bars = 250 µm.

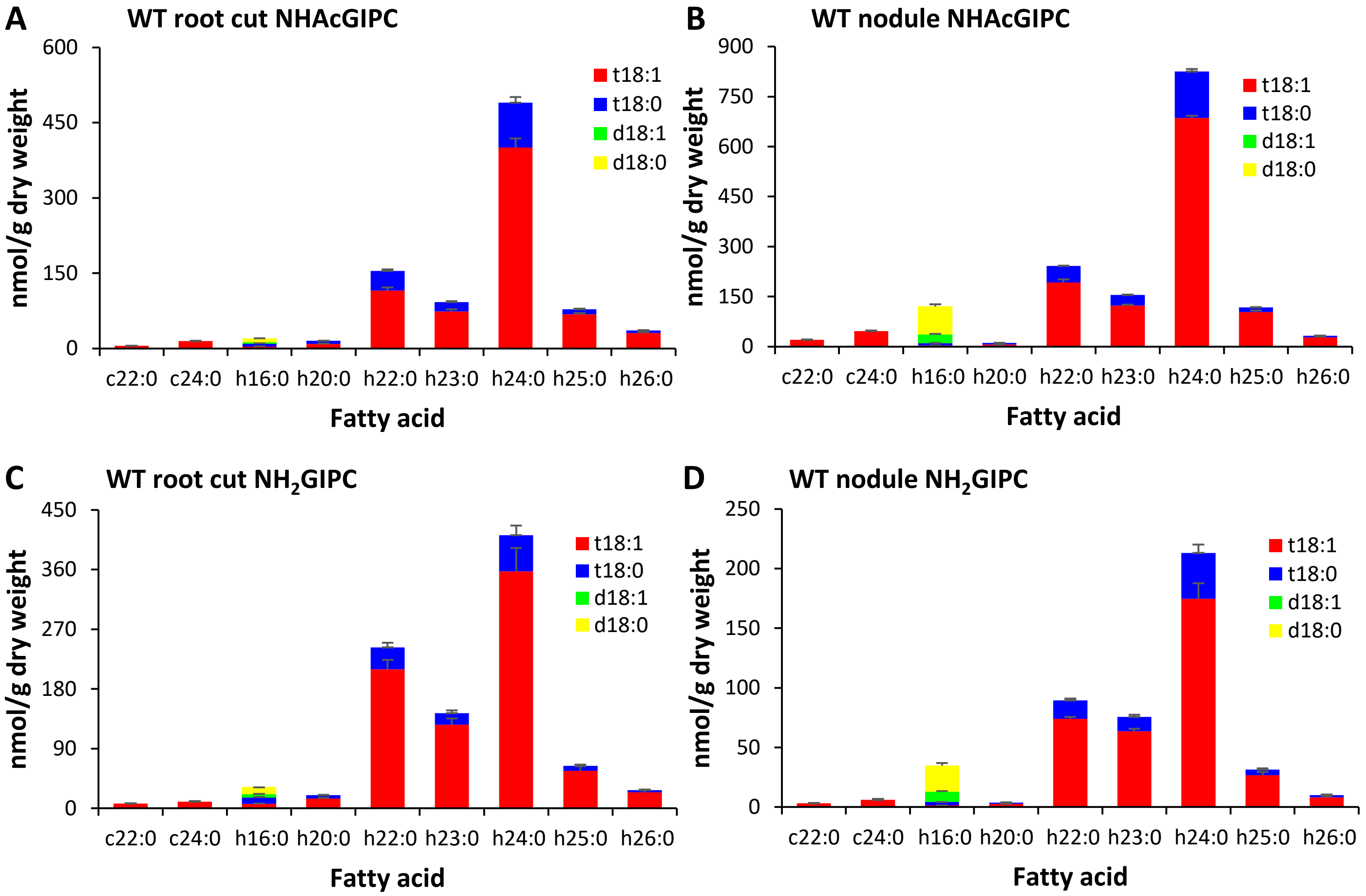


**Supplemental Figure S6.** Comparison of NHAcGIPC and $NH_2$GIPC molecular species composition between the WT roots and nodules after inoculation with *Bradyrhizobium* ORS278. (**A**) and (**B**) NHAcGIPC molecular species in WT roots after removal of nodules (**A**) and in nodules (**B**). (**C**) and (**D**) $NH_2$GIPC molecular species in WT roots after removal of nodules (**C**) and in nodules (**D**). Values are means ± SE from three technical replicates. The experiments were performed twice with similar results. The results of one representative experiment are shown.

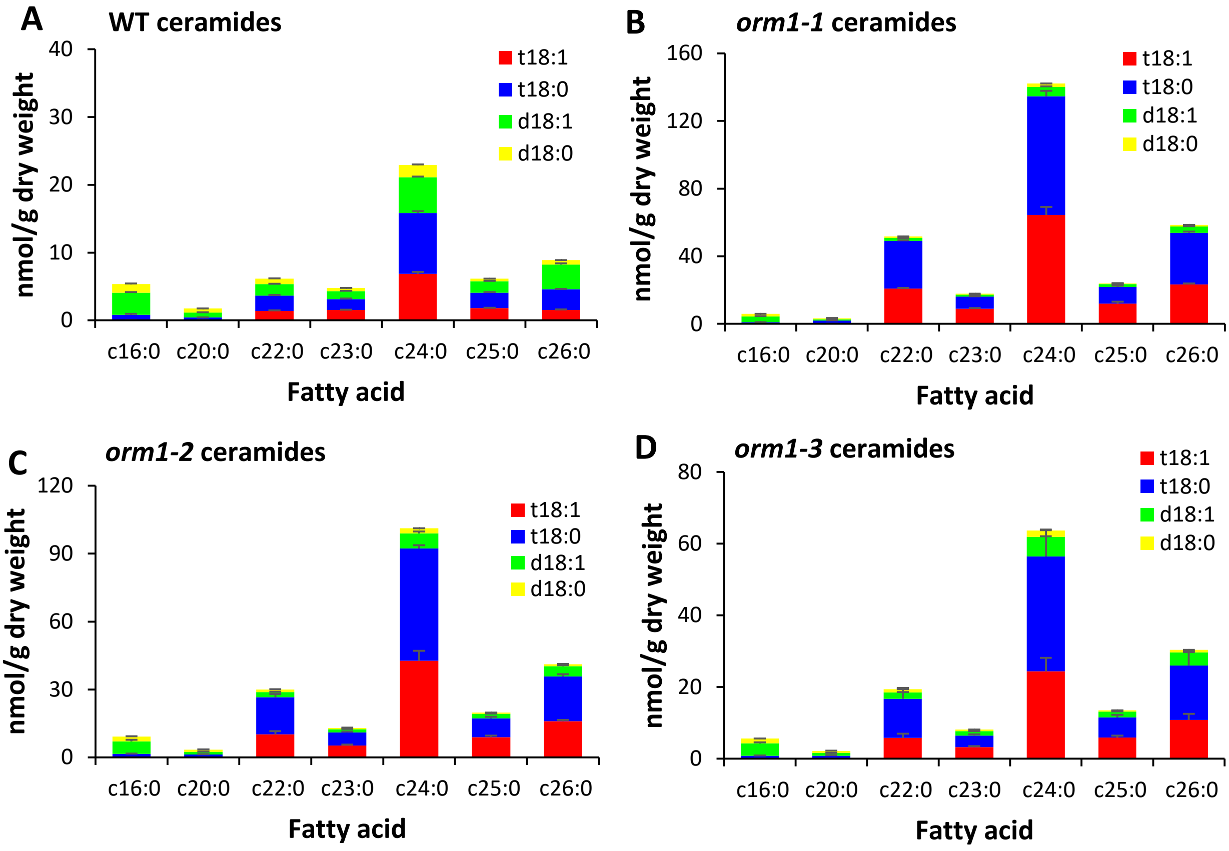


**Supplemental Figure S7.** Ceramide molecular species composition representing the exact pairings of LCB and fatty acid in roots of WT and *orm1* mutant plants. (**A**) to (**D**): (**A**) WT, (**B**) *orm1-1* mutant, (**C**) *orm1-2* mutant, and (**D**) *orm1-3* mutant. Values are means ± standard deviations (s.d.) from three technical replicates. The experiments were performed twice with similar results. The results of one representative experiment are shown.
