## Supplemental Tables 1-4 for "*ORM*-mediated regulation of sphingolipid biosynthesis is essential for nodule formation in *Aeschynomene evenia*"

**Supplemental Table S1.** Genetic data on the *Aeschynomene evenia orm1* mutants.

| | | | Mutation effect | | Genetic determinism | |
|---|---|---|---|---|---|---|
| **Gene** | **Mutant** | **Allele** | **Nucleotide change** | **Amino-acid change** | **F2 segregation WT:mutant** | **(*, P> 0,05)** |
| *AeORM1* (Ae02g01550) | P35 | *orm1-1* | c.224C>T | p.Pro75Leu | F2 489:158 | 3:1* |
| | Q33 | *orm1-2* | c.283G>A | p.Gly95Arg | F2 464:126 | 3:1* |
| | AG2 | *orm1-3* | c.74G>A | p.Gly25Asp | F2 458:96 | 3:1* |

**Supplemental Table S2.** Allelism analysis of the *Aeschynomene evenia orm1* mutants.

| Gene | Crossing (♀ x ♂) | n F1 plants (n pods) | F1 phenotype |
|---|---|---|---|
| *AeORM1* (Ae02g01550) | P35 x AG2 | 13 (2) | mutant |
| | Q33 x P35 | 22 (3) | mutant |
| | Q33 x AG2 | 11 (3) | mutant |

**Supplemental Table S3.** Complementation of the nodulation phenotype of *orm1-1* mutant plants by expressing WT *AeORM1* .

| ***A. evenia* mutant** | **Transformation construct** | **Transformed plants** | **Nodulated plants** | **Nodules/ nodulated plant*** |
|---|---|---|---|---|
| P35 | pCambia2200DsRED empty vector (EV) | 16 | 0 | 0 |
| P35 | pCambia2200DsRED-ProAeORM1:AeORM1 | 35 | 27 | 3.3±0.37 |

* values correspond to mean nodule number per nodulated plant ± standard deviation

**Supplemental Table S4.** Primer sequences used for qRT-PCR.

| Gene | Primer name | Sequence | Reference |
|---|---|---|---|
| *AeCRK* | AeCRK-F | CCCCAATAGTTCTGAGCCAACC | Quilbé et al. (2022) |
| | AeCRK-R | AGGAAGAGGCCATAGAGCCTTG | |
| *AeEF1a* | AeEF1-F | TGCTGGTATGGTTAAGATGGTTCC | Quilbé et al. (2022) |
| | AeEF1-R | TTCTTCTTCTGTGCTGCCTTGG | |
| *AeENOD40* | AeNOD40-F | CACACTTCTCCTCCATTCACTTTTC | Quilbé et al. (2022) |
| | AeNOD40-R | TTGCCATACTTGTAGCCAAAAGC | |
| *AeNIN* | AeNIN-F | CAACAGAACAAGGGGAAAGGGG | Quilbé et al. (2022) |
| | AeNIN-R | TAATGAGGCAGAGGCGGAAGTG | |
| *AeORM1* | AeORM1-F | AATGAGAATACCTCTTTTTGC | This study |
| | AeORM1-R | CCATATCCTTTCCAGTTCTGAATCC | |
| *AeORM2* | AeORM2-F | AACGATAGACCGATAACATATAC | This study |
| | AeORM2-R | GTATAGACTTCACAACTGGAAG | |
| *AeRAM1* | AeRAM1-F | GTAATTCCCTCGGTAATCTGTT | Quilbé et al. (2022) |
| | AeRAM1-R | TGGCTTCCTGCTCAACTA | |
| *AeSBT* | AeSBT-F | ATGAAGGAATGCCACCTCCACC | Quilbé et al. (2022) |
| | AeSBT-R | TGTGTGTGCCGTGTCCATTATC | |
| *AeSBTM1* | AeSBTM1-F | TGATATAGGCGGAGGAGT | Quilbé et al. (2022) |
| | AeSBTM1-R | ACATGAACAGCAGGAAGTA | |
| *AeSTR* | AeSTR-F | TTCCTCGTATGTCATATCTTCA | Quilbé et al. (2022) |
| | AeSTR-R | GCATTGGTTGTGATAAGTGA | |
| *AeSYMREM1* | AeSYMREM1-F | TGATGATGCTGCTGATGA | Quilbé et al. (2022) |
| | AeSYMREM1-R | TTTGGGTTGTGAAGAAGTG | |
| *AeVPY* | AeVPY-F | CTGAGGCTTCTCTTGCTTA | Quilbé et al. (2022) |
| | AeVPY-R | CTATGGCTGCTGCTATGT | |
| *AeUbi* | AeUbi-F | TCAAAGTGAAGACTCTAACCG | Quilbé et al. (2022) |
| | AeUbi-R | CAAGTGAAGCACGGAACC | |
| *RiGADPH* | RiGADPH-F | GACGTCTCAGTTGTTGATTTA | Buendia et al. (2016) |
| | RiGADPH-R | TTTGGCATCAAAAATACTAGA | |
| *RiLSU* | RiLSU-F | GCATATCAATAAGCGGAGGA | Xue et al. (2015) |
| | RiLSU-R | ACTCCTCACGCTCCACAGA | |
